# Intestinal control of feeding initiation in *Drosophila melanogaster*

**DOI:** 10.1101/2025.03.31.646269

**Authors:** Cindy Reinger, Laura Blackie, Hugo Gillet, Pedro Gaspar, Alexandra M. Medeiros, Dafni Hadjieconomou, Michèle Sickmann, Markus Affolter, Irene Miguel-Aliaga, Anissa Kempf, Martin Müller

## Abstract

The interplay between feeding and excretion is essential for organismal nutrition and survival, yet their mechanistic coupling remains poorly understood. At the onset of life, feeding must be initiated while developmental waste products – the meconium – need to be eliminated. Using Drosophila as a model system, we explored the in vivo mechanisms coordinating these processes. We developed novel behavioral assays for newly eclosed flies and discovered that, similar to neonatal mammals, Drosophila excrete their meconium shortly after eclosion. Remarkably, feeding initiation occurs only after partial meconium elimination. We identified a cis-regulatory element associated with the *apterous* gene, which, when disrupted, prevents both meconium excretion and adult feeding initiation. These flies develop hindgut obstruction (ileus), avoid food, and exhibit increased proboscis extension sleep – a behaviour we found plays a functional role in waste clearance under normal conditions. Through experimental inhibition of meconium excretion, we established that this process is prerequisite for feeding initiation, suggesting a gut-to-brain signaling circuit that couples these fundamental physiological processes. The progression of phenotypes we observed parallels the hallmarks of mechanical ileus in humans. Our findings reveal previously unrecognized links between intestinal clearance, feeding behavior, and survival, with potential implications for understanding similar processes across species.

## Introduction

The survival of animals hinges on a precisely orchestrated interplay of physiological processes. Among these, meconium excretion and feeding initiation represent crucial developmental milestones that establish the foundation for independent nutrient acquisition and growth across diverse taxa. Despite their apparent evolutionary conservation, the mechanisms coordinating these essential processes remain unexplored.

Meconium excretion marks a critical transition in early life. This process eliminates accumulated metabolic waste from embryonic or pupal development (in vertebrates and arthropods, respectively), manifesting as an odorless, mucilaginous substance whose colour varies from greenish-black to yellow-reddish across species. In mammals, failure to expel this developmental byproduct has been linked to severe conditions such as cystic fibrosis (Sathe and Houwen, 2017; Waldhausen and Richards, 2018), while its premature release can lead to meconium aspiration syndrome—a major cause of neonatal mortality (Ahanya et al., 2005).

Feeding initiation represents an equally pivotal milestone, triggering the first intake of external nutrients. This process activates metabolism, stimulates gut motility, and enables somatic growth – yet, how does the organism know when to begin feeding? Recent advances have illuminated the neurological underpinnings of feeding initiation in both vertebrates and arthropods (Betley et al., 2015; Chen et al., 2015; Youn et al., 2018, Miroschnikov et al., 2020; Xiao et al., 2022; Alcantara et al., 2022; Liu et al., 2023), but potential connections to excretory processes remain unexplored.

Correlative evidence across diverse species hints at an intimate interdependence between these two processes. Equine neonates with meconium retention show diminished feeding interest; myliobatiform stingrays eliminate meconium before initiating feeding; and human infants with meconium plug syndrome display difficulties in food intake (Rakestraw and Hardy, 2006; Tomita et al., 2020; Messina et al., 2016). Could meconium excretion be causal to feeding initiation? Do these observations reflect a conserved developmental strategy across animal phylogeny? How might the precise timing of these processes influence organismal health and survival?

*Drosophila melanogaster* is a genetically tractable model organism widely used for studying fundamental physiological processes. Despite their evolutionary divergence, *Drosophila* and vertebrates share fundamental structural and functional characteristics in their digestive systems, making flies an excellent model for investigating universal principles of intestinal physiology (Apidianakis and Rahme, 2011; Buchon et al., 2013; Miguel-Aliaga et al., 2018; Medina et al., 2022). In this study, we investigate a century-old uncharacterized phenotype in *Drosophila* mutants lacking the *apterous* (*ap*) gene (best known for its role in wing development; Metz, 1914). By combining modern genetic tools with behavioral assays, we reveal unexpected links between meconium excretion, hindgut function, sleep and feeding initiation.

## Results

### Meconium excretion and adult feeding initiation are temporally coupled

Upon eclosion, several organs of newly hatched flies undergo a series of developmental transitions to achieve full functional maturity, including the gut, the reproductive organs and the wings (Lemaitre and Miguel-Aliaga 2023; Hinnant et al 2020; Diaz de la Loza and Thompson 2017). The female reproductive organs, in particular the ovaries, remain underdeveloped at eclosion and require hormonal signals and nutrient availability to initiate vitellogenesis, leading to a dramatic increase in ovary size (Meiselman et al., 2018; reviewed in Bownes, 1982). Similarly, the gut, initially filled with meconium — waste products accumulated during metamorphosis — undergoes substantial remodeling and gradually adopts its adult structure over several days (O’Brien et al 2011; Marianes and Spradling 2013; Buchon et al., 2013). This post-eclosion remodeling seems to be tightly coordinated with the initiation of adult feeding, ensuring a smooth transition from metamorphosis to adult life.

To better understand the coordination between feeding and excretion, we examined the chronological relationship between meconium excretion and the initiation of adult feeding in wild-type flies (in this study, flies with a *y w* genetic background are used as wild-type; referred to as control (or ctrl) in the text). We found that meconium deposits, identifiable by their greenish coloration, were excreted in multiple steps over several hours (Fig. 1A, B). Despite sexually dimorphic traits in feeding and metabolism (Magwere et al 2004; Regan et al 2016), the dynamics of meconium excretion were comparable between males and females (Fig. 1B).

**Figure 1.**
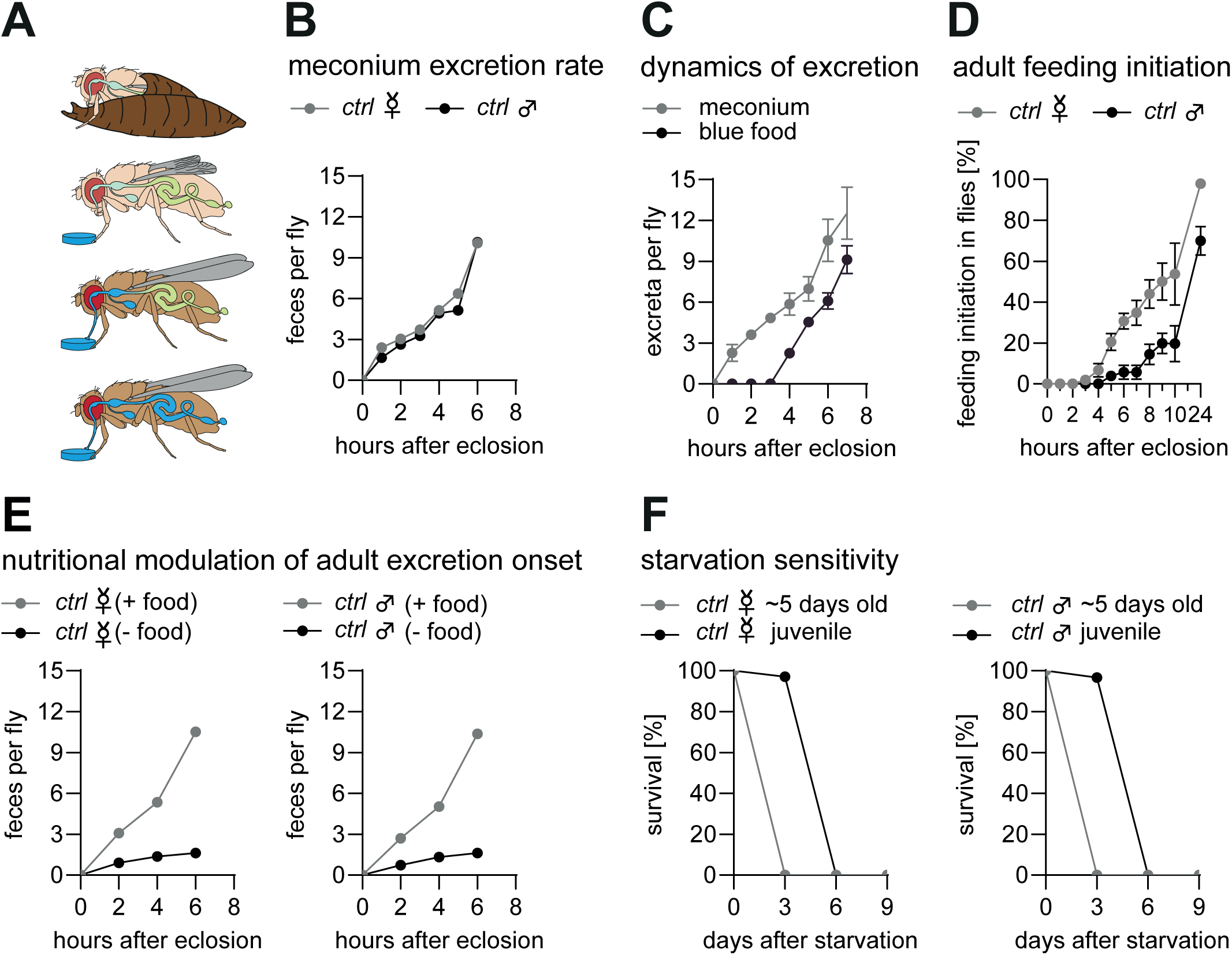
Meconium excretion is temporally coupled to feeding initiation. **A:** Schematic overview of feeding-related physiological events during the first 24 hours post-eclosion in *Drosophila melanogaster*. Chronological order is from top to bottom. Juvenile flies emerge from the pupal case. After eclosion, the meconium (green) is excreted in multiple steps. Adult feeding begins following partial meconium excretion, and digested food (blue) gradually fills the gut. Several hours post-eclosion, the intestine is entirely filled with food, and the meconium is fully expelled. **B.** Meconium excretion occurs in multiple steps over a period exceeding 6 hours in both sexes. Excretion begins within the first hour after eclosion. *n* = 25-30 flies per time point. **C.** The first digested food appears in excreta 4 hours post-eclosion. Each data point represents the average of n=12–19 samples pooled across four independent experiments. **D.** The first *ctrl* flies initiate adult feeding approximately 3 hours post-eclosion. Females begin feeding earlier than males, and by 24 hours, all females have ingested food, whereas only 70% of males have done so. Each data point represents the average of *n* = 12–19 pooled across four independent experiments. **E.** Meconium excretion rate is slower in the absence of food in both sexes. Each data point represents the average of *n* = 24-25 flies pooled across four independent experiments. **F.** Freshly eclosed *ctrl* flies exhibit enhanced survival under starvation conditions, living approximately three days longer than 5-day-old starved *ctrl* flies. Each data point represents the average of *n* = 25 pooled across three independent experiments. Data are means ± s.e.m. *n*, number of flies.

To examine when adult flies initiate feeding, freshly eclosed flies were kept in a petri dish with a piece of blue-dyed food. Ingested blue food and its excreta are readily discernible through its accumulation in the abdomen and as blue deposits in the petri dish, respectively. Freshly eclosed flies exposed to blue-dyed food exhibited visible ingestion approximately three hours post-eclosion (Fig 1D), with the first blue-dyed excreta appearing after four hours (Fig. 1C). Notably, females exhibited an earlier onset of feeding: 100% of females initiated feeding within 24 hours, whereas only ∼70% of males had done so (Fig. 1D). To assess how feeding initiation affects meconium excretion, freshly eclosed flies were kept in a petri dish without food, and meconium deposits were quantified. In the absence of food, meconium excretion was significantly delayed in both sexes, suggesting that a food-derived cue(s) facilitates the process (Fig. 1E).

Absence of food has other and seemingly more dramatic consequences for juvenile flies. For instance, oogenesis is suspended at a regulatory checkpoint that prevents the onset of vitellogenesis (Drummond-Barbosa and Spradling, 2001; reviewed in Schwenke et al., 2016). Additionally, the breakdown of energy-storing larval fat cells is significantly delayed by several days (Aguila et al., 2007). Without access to food, flies typically survive no longer than 2–3 days (Linford et al., 2013). To confirm this, we assessed the survival of freshly eclosed and well-fed five-day-old control male and female flies. Regardless of sex, five-day-old flies died within three days of starvation. Interestingly, despite never having consumed food, juvenile flies survived up to three days longer (Fig. 1F; see also Aguila et al., 2007). We have characterized a *Drosophila* mutation that, when kept on rich diet, elicits a death rate highly similar to starving control flies. We will show that in this mutation, meconium excretion is prevented. This observation served us as a means to compare and elucidate the distinct temporal sequence between meconium excretion and feeding initiation in control and mutant flies.

### Meconium retention induces a starvation-like phenotype in *ap* mutant flies

Among thousands of *Drosophila* mutations known so far, *ap* mutants exhibit a survival pattern that closely resembles starvation-induced lethality in wild-type flies (Metz 1914). The *ap* gene codes for a LIM-homeodomain transcription factor (Cohen et al., 1992) and is best known for its role as a dorsal selector gene during wing formation, where it establishes dorso-ventral compartmentalization in imaginal wing discs (Butterworth and King, 1965; Stevens and Bryant, 1985, 1986; Cohen et al., 1992; Milán and Cohen 2000; Bieli et al., 2015a,b; Aguilar et al 2023). Additionally, Ap plays a role in the development of the embryonic and larval nervous system (Cohen et al., 1992; Lundgren et al., 1995; O’Keefe et al., 1998; Herzig et al., 2001) as well as in the development of embryonic muscle (Capovilla et al., 2001). However, very little is known about its function to enable an adult fly to survive longer than three to four days.

Through detailed genetic analysis of the *ap* locus, we identified a putative regulatory DNA element referred to as the Life Span Enhancer (LSE) (Fig. 2A, Supp. Fig. 1A), which is required and sufficient to promote adult survival (Reinger et al, manuscript in preparation). We found that over 80% of homozygous *ap* mutant flies, which lack both copies of the LSE but carry intact copies of the *ap* coding region (referred to as *ap^ΔLSE^*flies; Fig. 2A, Supp. Fig. 1D), die within 3 days post-eclosion as opposed to heterozygous *ap^ΔLSE^/+* and controls (Fig. 2A,B). Reintroducing one copy of the LSE into an LSE-deleted background was sufficient to fully rescue the survival phenotype (referred to as *ap^minLSE^*; Fig. 2B, Supp. Fig. 1G). This shows that the loss of the LSE is responsible for the survival phenotype, and that *ap^ΔLSE^* mutant flies grown on rich medium exhibited mortality rates similar to those of starved control flies (cf. Fig. 1F).

**Figure 2.**
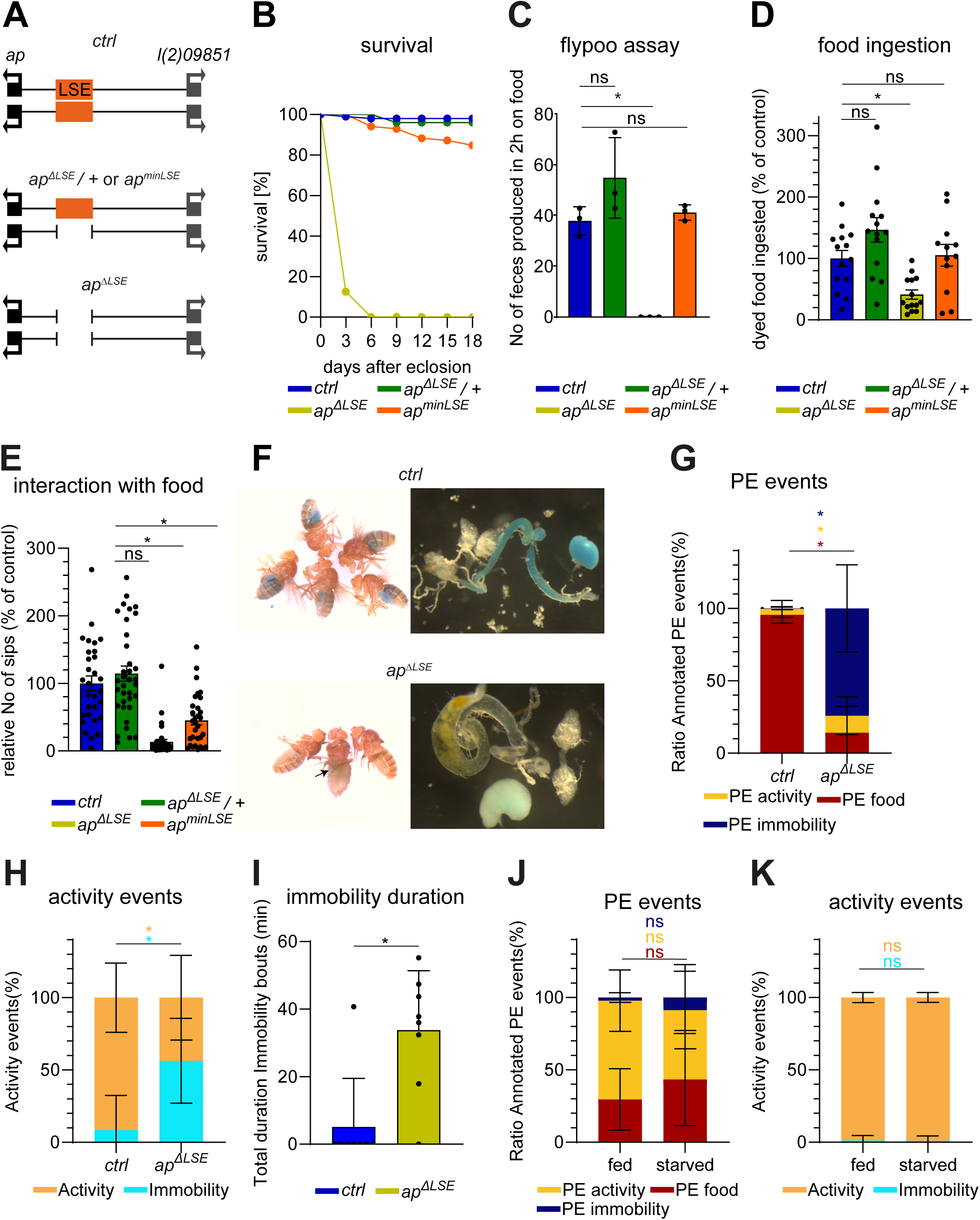
The *apterous* related Life Span Enhancer rescues meconium excretion, feeding initiation and survival of adult flies. **A.** Schematic representation of the *ap* locus. *ap* is depicted in black and its flanking gene *l(2)09851* to the right is depicted in grey. The LSE is depicted in orange. Top: wild-type *ap* locus in which the LSE is located in the intergenic spacer. Middle: heterozygous flies with the LSE present on one chromosome and deleted on the other. Bottom: deletion of the LSE in the *ap^ΔLSE^* mutant, resulting in loss of regulatory activity. **B.** Survival curves of different *apterous* (*ap*) alleles. In the presence of the life span enhancer, more than 80% of flies survive 18 days at 25°C. In contrast, flies lacking the LSE exhibit premature mortality similar to starved control flies. This phenomenon is referred to as precocious adult death. Data are based on *n* = 100-120 flies. **C.** Flypoo assay: bar graphs depict the total number of fecal deposits of 1 day old flies maintained on blue-dyed food over a period of 2 hours. *ap^ΔLSE^*flies show no detectable fecal output, indicating severe intestinal blockage. Each data point represents the average of *n* = 30 pooled across three independent experiments. **D.** Bluefood squash assay: The mean amount of ingested dye per fly over 1 hour of feeding is reduced in one-day-old *ap^ΔLSE^* female virgins. Flies were starved overnight and put on blue-dyed food the day after. Each data point represents the average of *n* = 8-16 pooled across three independent experiments. **E.** FlyPad assay: the relative number of proboscis sips per fly over 1 hour of feeding is reduced in one-day-old *ap^ΔLSE^* female virgins. Flies were kept under *ad libitum* conditions after eclosion. Each data point represents the average of *n* =8-12 pooled across three independent experiments. **F.** Gut content analysis in one-day-old flies. Top: images of one-day old *ctrl* flies constantly kept on blue food. The blue dye can be detected in crop and midgut. Bottom: images of one-day-old *ap^ΔLSE^* flies constantly kept on blue food. In rare cases, blue dye can be detected in the crop (arrow). Note that the midgut is devoid of blue color and contains meconium (yellowish material in midgut on the left). **G.** Human annotation focused on 3 types of proboscis extensions (PE) with the following criteria: proboscis extended on food (red); proboscis extended while moving (e.g., grooming), yellow); proboscis extended while remaining immobile (blue). Annotated data ratios show that *ctrl* flies direct most of their PE on food, while *ap^ΔLSE^* flies do so significantly more while immobile. *n* = 8. **H.** *ctrl* flies spend more time active than immobile. *ap^ΔLSE^* flies spend more time immobile than active. *n* = 8. **I.** The total duration of the immobility bouts shows that *ap^ΔLSE^*flies are sleeping. *n* = 8. **J.** Annotated data ratios show that *ctrl* and starved flies direct their PE equally on the three criteria. *n* = 8-12. **K.** *ctrl* and starved flies spend almost the same amount of their time active and immobile. *n* = 8-12. Asterisks indicate significant differences (*P*<0.05), ns = not significant (P>0.05). Data are means ± s.e.m. *n*, number of flies. See Supplementary Information for exact *P* values and statistical tests.

Further experiments revealed that *ap^ΔLSE^* flies were unable to excrete their meconium and to defecate as opposed to control, heterozygous and LSE rescue flies (Fig. 2C). Based on our findings regarding the temporal coupling of meconium excretion with feeding initiation (cf. Fig. 1), we hypothesized that *ap^ΔLSE^* flies may be unable to properly initiate feeding. Indeed, one-day-old *ap^ΔLSE^* flies barely ingested any food, with only occasional traces of blue-dyed food observed in the crop but never in the midgut (Fig. 2D, F, Supp. Fig. 2). Moreover, *ap^ΔLSE^* flies also displayed a significant reduction in food interaction events when compared to control and *ap^minLSE^* rescue flies (Fig. 2E), suggesting that their feeding behavior was severely impaired. To investigate whether the feeding phenotype of *ap^ΔLSE^* flies might be related to meconium retention, we dissected the midguts of control and mutant flies and assessed their contents. Dissections revealed that *ap^ΔLSE^*flies retained their meconium in their midguts (Fig. 2F) far beyond the expected time of excretion (cf. Fig. 1), supporting a link between meconium retention and feeding impairment.

To assess why one-day-old *ap^ΔLSE^* flies do not eat, we conducted high-resolution imaging analyses of *ctrl* and *ap^ΔLSE^*flies, and quantified their movement as well as different types of proboscis extension (PE) events over the one-hour experimental period: PE directed toward food, PE during immobility, and PE during active movement. In control flies, more than 90% of PE events were associated with food consumption, with only a minor proportion occurring during either locomotion, grooming or immobility. In contrast, *ap^ΔLSE^* mutants exhibited markedly reduced activity and predominantly displayed PE during immobility (∼80% of total PE events) (Fig. 2G, 2H, Supp. Fig. 3A).

So far, our observations indicate that in control flies, meconium is gradually replaced by ingested food during the first day of life. This suggests a distinct temporal sequence between meconium excretion and feeding initiation. The progression of events is severely disturbed in *ap^ΔLSE^* mutants because they are unable to excrete their meconium. This has conspicuous effects on their behaviour. They avoid food and become increasingly lethargic. Our video analyses indicated that proboscis dysfunction does not contribute to the observed feeding impairment. Rather, pronounced differences in the distribution of PE events were noted between control and mutant flies. They are described in the next section.

### *ap* mutant flies are engaged in proboscis extension sleep (PES)

The periodic occurrence of PE movements during immobility is highly reminiscent of proboscis extension sleep (PES), a distinct deep sleep state previously described in *Drosophila* (van Alphen et al., 2021). Unlike stereotypical feeding-associated PE, PES is characterized by unique dynamics and kinematics (van Alphen et al., 2021; Keles et al., 2025). Comparative analyses confirmed that the PE events displayed by *ap^ΔLSE^* mutants during immobility closely matched the defining features of PES (Supp. Fig. 4). Additionally, *ap^ΔLSE^* flies exhibited predominantly prolonged immobility bouts of ≥5 min inactivity – an established criteria of sleep in *Drosophila* (Shaw et al., 2000) –, further supporting that *ap^ΔLSE^* mutants are engaged in PES (Fig. 2I, Supp. Fig. 3B).

To determine whether the PES-like phenotype in *ap^ΔLSE^* flies could be attributed to a starvation-like state, we performed the same assay on one-day-old control flies subjected to acute starvation immediately post-eclosion. Neither fed nor starved control flies showed differences in the distribution of PE events across conditions (Fig. 2J, Suppl. Fig. 5A) and remained highly active throughout the experiment (Fig. 2K, Suppl. Fig. 5B).

These findings demonstrate that 1 day old *ap^ΔLSE^* flies are predominantly engaged in a PES state, and exclude starvation or energy depletion as underlying causes of the observed phenotype. Together with our observation that *ap^ΔLSE^* mutants fail to excrete their meconium, these results raise the intriguing possibility that the inability to eject meconium itself may sustain the prolonged sleep-like state observed in these mutants.

### The LSE mediates *ap* expression in the posterior hindgut

To understand how the LSE deletion triggered the feeding, excretion, and sleep phenotypes, we next investigated how and where this deletion affected *ap* expression. Ap is dynamically expressed throughout development in various tissues, for which several tissue-specific enhancers have been identified (Cohen et al., 1992; Capovilla et al., 2001; Herzig et al., 2001; Bergman et al., 2002; Lundgren et al., 1995; Gohl et al 2008; de Taffin et al., 2015; Bieli et al., 2015a,b; Stratmann and Thor, 2017). We hypothesized that our genetically characterized LSE corresponds to a tissue-specific regulatory element that activates the gene in a subset of Ap-expressing cells. To determine the anatomical site of action of LSE, we engineered enhancer-trap-like Gal4 driver lines in the *ap* locus with (*ap^Gal4^* and *ap^LSE-Gal4^*) or without (*ap^ΔLSE-Gal4^*) regulation by LSE, and traced their cell lineage using the G-TRACE system (Supp. Fig. 1H,I,J; for details on these alleles, see materials and methods). G-TRACE tracks all cells that currently express Gal4 (*ap*_current_) or have done so in the past (*ap*_cell lineage_) when activated by a Gal4 driver (Evans et al., 2009). Notably, the analysis of the Ap expression patterns immediately revealed where LSE is active: the posterior hindgut (Fig. 3). This finding is of significant interest given the observed feeding and excretion phenotypes of *ap* mutants.

**Figure 3.**
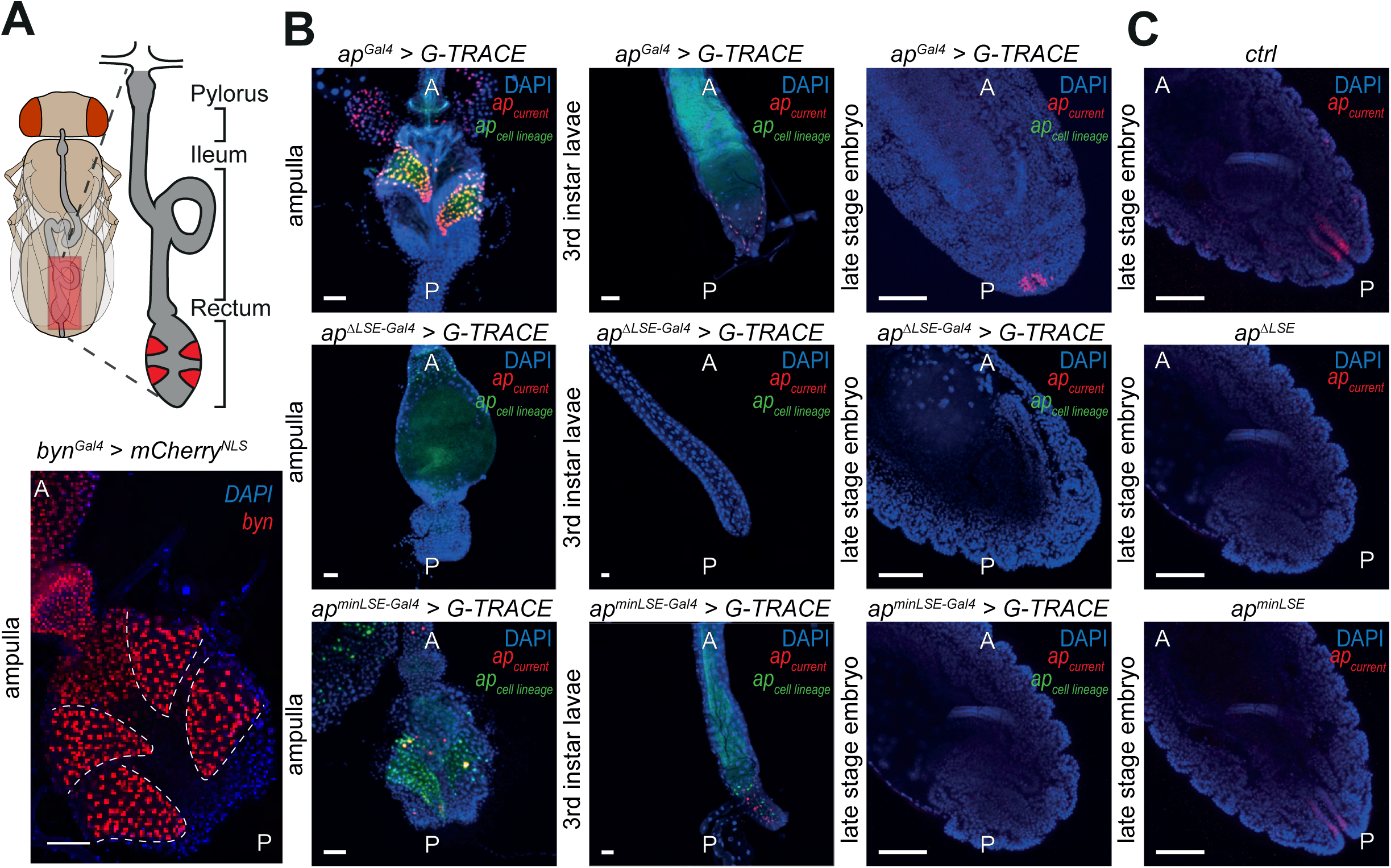
The *apterous* related life span enhancer is continuously expressed in the posterior hindgut during all developmental stages of the fly. **A.** Top: illustration of the posterior hindgut of adult *Drosophila*. Rectal papillae (red), responsible for reabsorption processes of ingested food, are located in the rectum. Bottom: the gene *brachyenteron* (*byn)* serves as a robust marker for rectal papillae exhibiting expression in all four papillae. **B.** G-TRACE fluorescence analysis of *apterous (ap) Gal4* driver lines focusing on the posterior hindgut across developmental stages. Fluorescence images reveal *ap* expression in the rectal papillae and hindgut cells during adult, larval and embryonic stages in *ap^Gal4^* and *ap^minLSE^* (one copy of the LSE) flies. If the LSE is absent on both chromosomes (*ap^ΔLSE^*), *ap* expression is absent in the hindgut across all stages. This confirms that the LSE is required and sufficient for hindgut specific *ap* expression throughout development. Blue: DAPI. Red: current expression of *ap*. Green: cell lineage expression of *ap*. **C.** Confirmation of tissue specificity of *ap^Gal4^* driver lines by antibody stainings of posterior hindguts in late stage embryos. As shown in **B**, *ap* is not expressed in the hindgut of *ap^ΔLSE^* embryos. Presence of the LSE rescues *ap* expression in the hindgut. Blue: DAPI. Red: Ap detected with anti-ap antibody. A = anterior, P = posterior. Scale bar = 50 μm.

The hindgut consists of three subdivisions: pylorus, ileum and rectum (Fig. 3A; reviewed in Cohen et al., 2020). A key structure within the adult rectum is the ampulla, which contains four cone-shaped rectal papillae that can be visualized using the *brachyenteron* (*byn*; Murakami et al., 1995) Gal4 driver (*byn^Gal4^*; Iwaki and Lengyel, 2002; Fig. 3A). Together with the Malpighian tubules, rectal papillae function analogously to mammalian kidneys (reviewed in Cohen et al 2020). We found that LSE-driven *ap* expression was restricted to two of the four rectal papillae and that this signal was completely absent in the hindguts of LSE-deleted (*ap^ΔLSE-Gal4^*) flies (Fig. 3B). Notably, our developmental G-TRACE analysis further revealed that *ap* was already expressed in the posterior hindgut of the embryo, where it labels two stripes that are absent in *ap^ΔLSE-Gal4^* flies (Fig. 3B, Supp. Video 1). Immunostaining of Ap in *ctrl*, *ap^ΔLSE^*, and *ap^minLSE^*embryos confirmed the G-TRACE expression patterns (Fig. 3C), validating the newly established Gal4 drivers as reliable reporters of Ap expression.

These findings show that LSE activates *ap* expression in the posterior hindgut as early as embryonic stages, and that *ap* expression persists in two of the four rectal papillae in adult flies.

### An ileus-like physical obstruction is present in the hindgut of *ap^ΔLSE^* flies

To determine how the absence of *ap* expression in the hindgut of *ap^ΔLSE^* mutants affects rectal papillae formation, we examined dissected ampullae from one-day-old adult *ap^ΔLSE^* flies using widefield and confocal imaging. In control flies, four papillae per ampulla are formed - each targeted by an intricate tracheal network (Fig. 4A; Wessing and Eichelberg, 1973) - and express the papilla marker *Delta* (*Dl*) (Fig. 4A’). In *ap^ΔLSE^* flies, papillae are absent, and instead, a malformed structure targeted by a tracheal jumble forms at the posterior end of the ampulla, which we term the Reinger’s knot (Fig. 4B,B’). This structure appears to consist of mispositioned papilla cells that failed to separate and properly integrate in the ampulla, as indicated by persistent Dl expression (Fig. 4B’). These imaging experiments show that the rectal papillae of adult *ap^ΔLSE^* flies fail to form properly, and also suggest that the Reinger’s knot fully obstructs the hindgut, potentially explaining why *ap^ΔLSE^* flies are unable to excrete their meconium. Note that the hindgut posterior to the Reinger’s knot is not obstructed in *ap^ΔLSE^*flies (Supp. Fig. 6). This occlusion functionally resembles an ileus in humans, a condition that, if untreated, is lethal (Plusczyk et al. 2006; Ansari, 2024; Wang et al., 2024).

**Figure 4.**
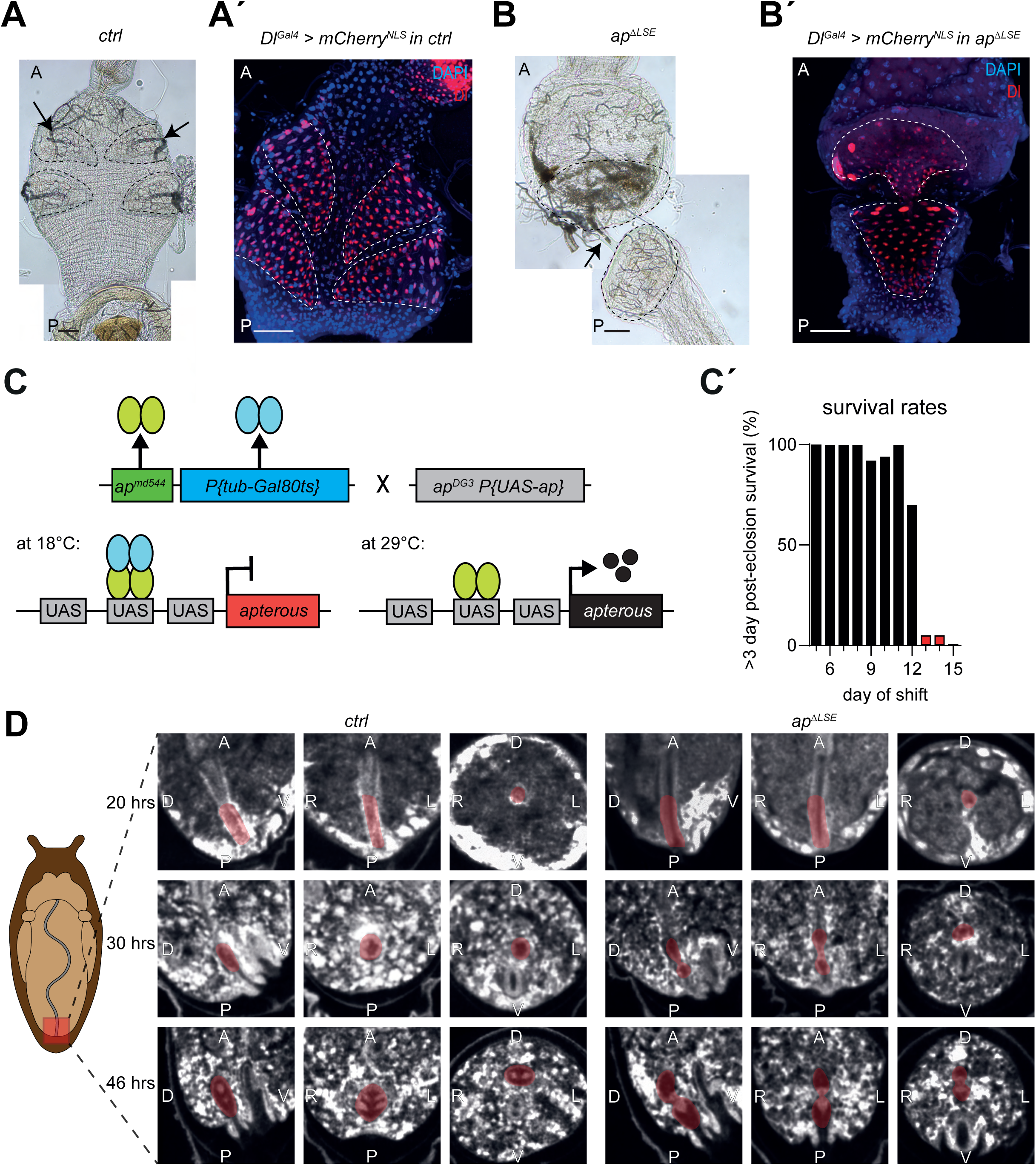
The hindgut-obstructing Reinger’s Knot is formed in the absence of the *apterous* related LSE. **A.** The rectal papillae (outlined by black dotted lines) require a substantial oxygen supply to perform their functions in reabsorption and waste clearance. Therefore, they are innervated by trachea (arrows). Each papilla contains a well-organized tracheal network in adult *ctrl* flies. **Á.** *Delta* (*Dl*), a cell surface protein, is present in rectal papillae cells. The mCherry^NLS^ signal (red) in *Dl^Gal4^ ctrl* flies highlights all four rectal papillae (outlined by white dotted lines). **B.** The Reingeŕs knot (outlined by black dotted lines) consists of an anterior and a posterior part connected by a small duct (black arrow). Meconium accumulates in the anterior part of the Reinger’s knot and is visible as a brownish substance within the duct. The knot is targeted by an unorganized trachea network. **B′.** Reingeŕs knot (outlined by a white dotted line) also expresses *Dl* (red) in an *ap* mutant background. All observations indicate that the knot formed in *ap^ΔLSE^* flies consists of papilla derived tissue. **C.** *tubGal80ts*-based temperature-shift experiments were performed to determine the period of *apterous* (*ap*) function relevant for proper papillae formation. *ap^md544^,* a *bona fide ap*^Gal4^ driver line and a strong *ap* allele (green), was recombined with *tubGal80^ts^* (blue), and crossed with *ap^DG3^ P{UAS-ap}*. *ap^DG3^* lacks the LSE (grey). At 18°C, Gal80^ts^ inhibits Gal4 produced by *ap^md544^*and no Ap protein can is synthesized. At 29°C, Gal80^ts^ is inactive and Ap protein can now be produced under the control of Gal4. **C’.** Animals shifted to 29°C (and therefore expressing Ap) before day 13 survived normally (black bars), whereas those shifted after day 12 displayed the precocious adult death phenotype (red bars). This indicates that a 24 hrs pulse of Ap expression in early metamorphosis is sufficient to ensure survival of the flies. *n* = 10 – 20 (see Supp. Table 3). **D.** Left: Schematic overview of a *Drosophila* pupa (light brown) within its pupal case (dark brown). The intestine is depicted in grey and the location of the posterior hindgut with rectal ampulla is highlighted in red. Right: µ**-**CT scans of papillae formation in *ctrl* and *ap^ΔLSE^* pupae at 20, 30 and 46 hours apf. At 20, no differences in the ampulla (highlighted in red) is observed in both conditions. At 30 hours apf, in *ctrl* flies, two regions of papilla tissue become visible, positioned dorsally and ventrally around the ampulla. In *ap^ΔLSE^* flies, however, a bulbous tissue structure is present centrally within the ampulla. At 46 hours apf, four papillae are formed in the ampulla of *ctrl* flies. In *ap^ΔLSE^*flies, papillae are malformed. Tissue remains centrally positioned and does not form four papillae. This newly formed structure is called the Reinger’s knot. Scale bar = 50 μm. A=anterior. P=posterior. D=dorsal. V=ventral. L=left. R=right.

To identify the developmental stage at which *ap* is required for rectal papillae formation, we performed temperature-sensitive rescue experiments using the Gal4/Gal80ts/UAS system to restore *ap* expression into an otherwise mutant background (Fig. 4C). We found that transient expression in the hindgut during the first day of metamorphosis is sufficient to rescue precocious lethality (Fig. 4C, C’; Supp. Table 3; McGuire et al., 2004), suggesting that *ap* may play a key role in rectal papillae formation during early metamorphosis (see also Wilson, 1981a).

To test this, we performed µ-CT imaging (Schoborg et al, 2019; Blackie et al 2024) on 20, 30 and 46 hours-old control and *ap^ΔLSE^* pupae to visualize internal structures throughout development (Fig. 4D). Differences in hindgut morphology first appeared at 30 hours after pupa formation (apf) with rectal papillae forming at the ampulla’s edges of control flies, while remaining centrally positioned in *ap^ΔLSE^* mutants. By 46 hours apf, control pupae had four distinct papillae, whereas *ap^ΔLSE^* mutants retained a central mass obstructing the hindgut. This suggests that papillae formation is arrested at the stage when the rectal tube normally splits into four cone-shaped luminal structures (Schoenfelder et al 2014).

The 3D visualization of the entire gut of one-day-old control, *ap^ΔLSE^* and *ap^minLSE^* flies revealed an additional phenotype in the midgut. While control and *ap^minLSE^* midguts appeared normal, it was bloated in *ap^ΔLSE^* flies (Supp. Fig. 7A). Actin immunostaining showed no structural abnormalities in the visceral mesoderm of *ap^ΔLSE^* flies (Supp. Fig. 7C), but by day two, *ap^ΔLSE^* midguts frequently showed signs of decay (Supp. Fig. 7D), suggesting rupture and leakage into the abdominal cavity. This may explain the progressive darkening of the ventral abdominal epidermis observed in mutants (our unpublished observation).

In summary, our analysis of adult gut morphology in control and *ap^ΔLSE^* flies reveals the formation of the Reinger’s knot during early metamorphosis and subsequent midgut deterioration in 2-day old *ap^ΔLSE^* adults.

### Removal of the Reinger’s knot restores survival in *ap^ΔLSE^* flies

To assess whether the Reinger’s knot and/or midgut deterioration is responsible for the precocious adult death phenotype, we sought to prevent hindgut obstruction in *ap^ΔLSE^* flies by eliminating the Reinger’s knot. Since we found that the Reinger’s knot consists of mislocalized Dl-expressing papilla cells, we hypothesized that its removal could be achieved by eliminating these cells. A study on hindgut proliferation suggested that expressing the repressor isoform of *cubitus interruptus* (*ci^Rep^*; Aza-Blanc et al 1997; Müller and Basler, 2000) in the posterior hindgut using *byn^Gal4^* might prevent rectal papillae formation (see Fig. 4C in Takashima et al., 2008). We replicated this experiment and confirmed that *byn^Gal4^>ci^Rep^*flies completely lacked rectal papillae (Fig. 5A-A’’’), providing a means to ablate these cells. Notably, these flies remained viable under standard lab conditions but perished within nine days when raised on high-salt medium (250 mM; Fig. 5C, Supp. Fig. 8A).

**Figure 5.**
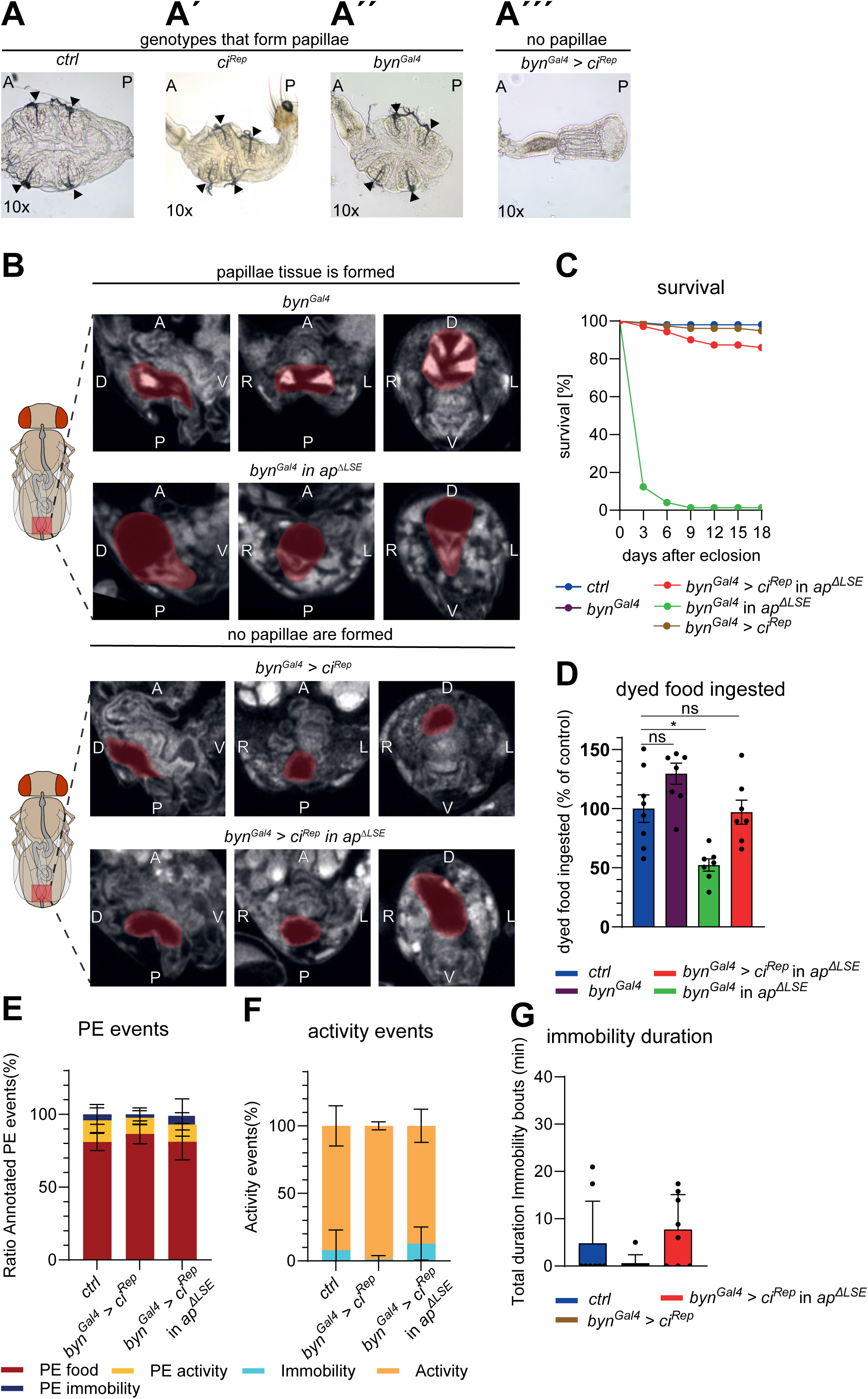
The removal of the Reingeŕs knot rescues excretion, feeding and consequently survival of *ap^ΔLSE^*flies. Ampullae of different genotypes were examined. A-A’’: In *ctrl*, *ci^Rep^* and *byn^Gal4^* flies, four papillae are located within the rectal ampulla (black arrow heads). A’’’: In *byn^Gal4^>ci^Rep^* flies, papilla tissue is absent. **B.** Top panel: on the left, a schematic overview of an adult fly indicates the position of the ampulla (highlighted in red) within the intestine (grey). On the right, µ-CT scans show that *byn^Gal4^/+* flies form rectal papillae (highlighted in red). The Reinger’s knot (highlighted in red) remains in place when *byn^Gal4^/+* is present in a *ap^ΔLSE^* mutant background. Bottom panel: papilla tissue is absent in a control as well as in a *ap^ΔLSE^* mutant background, because *byn^Gal4^*>*ci^Rep^* eliminates the formation of normal papillae as well as the Reinger’s knot, respectively. **C.** *T*he precocious adult death phenotype of *ap^ΔLSE^* flies is rescued upon removal of the Reinger’s knot by *byn^Gal4^>ci^Rep^*treatment. However, because papillae play an essential role in absorption processes, the rescue is not effective under high salt conditions (see Supp. Fig. 8A). n = 100 **D.** Feeding initiation can be rescued by *byn^Gal4^>ci^Rep^ in ap^ΔLSE^* flies. **E.** Like starved *ctrl* and *byn^Gal4^>ci^Rep^* flies, annotated PE ratios show that *byn^Gal4^>ci^Rep^ in ap^ΔLSE^*flies direct their PE equally distributed on all three criteria*. n = 8.* **F.** Activity events can be rescued in *byn^Gal4^>ci^Rep^ in ap^ΔLSE^*flies. *n* = 8. **G.** The sleep phenotype can be rescued in *byn^Gal4^>ci^Rep^* in *ap^ΔLSE^* flies. *n* = 8. A=anterior. P=posterior. D=dorsal. V=ventral. L=left. R=right. Asterisks indicate significant differences (*P*<0.05), ns = not significant (P>0.05). Data are means ± s.e.m. *n*, number of flies. See Supplementary Information for exact *P* values and statistical tests.

To determine whether *ci^Rep^* expression in the hindgut disrupts rectal papillae formation during metamorphosis similar to *ap*, we performed temperature-shift experiments using the *tub-Gal80ts* system (McGuire et al., 2004). Our findings indicate that *ci^Rep^* acts at the onset of metamorphosis, approximately one day before *ap* is required (Supp. Fig. 8B-B′′, Supp. Table 4; see legends for details).

To evaluate the effect of papilla removal on Reinger’s knot formation, we expressed *ci^Rep^* under *byn^Gal4^* control in *ap^ΔLSE^* mutants. µ-CT imaging confirmed the absence of rectal papillae and the elimination of the Reinger’s knot (Fig. 5B), demonstrating that *byn^Gal4^>ci^Rep^* disrupts papillae formation upstream of *ap*. µ-CT scans also showed that the midguts of the knot-ablated *ap^ΔLSE^* mutants were normal (Supp. Fig. 7B), strongly indicating that the Reinger’s knot is the primary cause of the bloated appearance and subsequent tissue decay observed in the midguts of *ap^ΔLSE^* flies (Supp. Fig. 7D). Notably, knot-ablated *ap^ΔLSE^* mutants are viable (Fig. 5C) and fertile. They also exhibited restored feeding behavior (Fig. 5D), activity levels (Fig. 5F,G, Supp. Fig. 9B) and showed no signs of PES (Fig. 5E, Suppl. Fig 9A) when compared to control flies.

Together, these findings show that the hindgut blockade (Reinger’s knot) observed in *ap^ΔLSE^*mutants is solely responsible for the survival, feeding and activity/sleep phenotypes, and underscores the critical role of *ap* in maintaining normal behavior in adult flies.

### Meconium retention by external obstruction induces precocious adult death

If meconium excretion is required for feeding initiation, then artificially blocking the anus of newly eclosed flies should produce opposite effects and mimic the *ap^ΔLSE^* phenotype. To test this, we applied superglue to the anus or to the anterior abdomen as a control (Fig. 6A). While application of super glue to the anterior abdomen had no adverse effects (Fig. 6B), obstructing the anus induced symptoms closely resembling those of *ap^ΔLSE^* mutants. Anus-sealed flies exhibited a sharp decline in survival, with approximately 80% perishing within three days, regardless of sex (Fig. 6B). Feeding was also impaired, and blue-dyed food was never detected in the midgut but occasionally in the crop (Fig. 6C, Supp. Fig. 2). Additionally, flies developed a brownish abdomen within one to two days, similar to *ap^ΔLSE^* mutants (unpublished data). Anus-sealed flies also showed a significant reduction in activity (Fig. 6E), and spent most of their time in PES (Fig. 6D, F, Supp. Fig. 10A, B).

**Figure 6.**
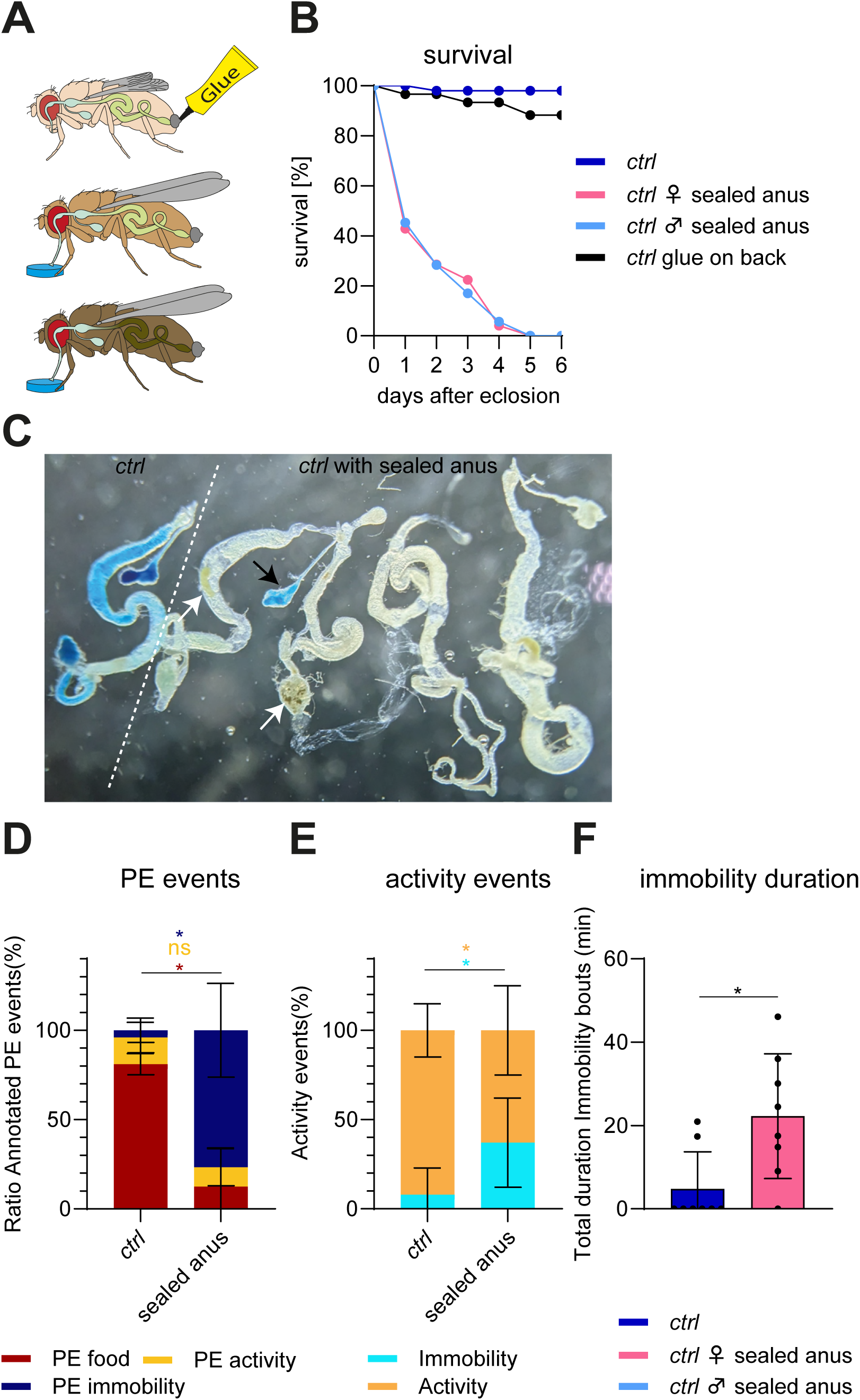
Flies with a sealed anus phenocopy the behaviour of *ap^ΔLSE^* mutant flies. **A.** Schematic overview of the sealed-anus experiment in chronological order from top to bottom. Freshly eclosed flies were shortly anaesthetised and a drop of super glue was applied to their anus to prevent meconium excretion. As a result, flies with a sealed anus did not initiate feeding, and as observed for *ap^ΔLSE^* flies, developed a darker appearance. **B.** The induced constipation phenotype caused by sealing the anus resulted in mortality dynamics similar to those observed in *ap^ΔLSE^* flies. Both *ctrl* groups survived normally. *n* = 50. *n* = 25 for *ctrl* with glue on the back. **C.** Like in *ap^ΔLSE^* flies, *ctrl* flies with a sealed anus failed to initiate adult feeding indicated by the absence of ingested blue diet in dissected 24 hours old intestines. In rare cases, food enters the crop (black arrow) but never the midgut. Meconium leftovers (white arrows) can be detected within the intestine. **D.** Annotated PE ratios show that *ctrl* flies direct most of their PE on food, while constipated flies do significantly less so, and show more PE when immobile. *n* = 8. **E.** As opposed to *ctrl* flies, constipated flies spend significantly more of their time immobile. *n* = 8. **F.** The total duration of immobility bouts shows that constipated flies are sleeping. *n* = 8. Asterisks indicate significant differences (*P*<0.05), ns = not significant (P>0.05). Data are means ± s.e.m. *n*, number of flies. See Supplementary Information for exact *P* values and statistical tests.

Together, these results show that meconium retention by external obstruction is sufficient to prevent feeding initiation, and to induce a PES state as well as precocious adult death. These findings highlight a previously unrecognized link between meconium excretion, feeding initiation and survival.

## Discussion

Our study identifies a previously uncharacterized developmental and physiological process in *Drosophila melanogaster*: the temporal coupling between meconium excretion and the initiation of adult feeding. We show that the successful transition to post-eclosion feeding depends on the partial or complete expulsion of the meconium suggesting a physiological checkpoint that coordinates these events. Additionally, we provide a genetic framework underlying the meconium-induced constipation of *ap^ΔLSE^* flies, which results in a starvation-like, lethargic phenotype coupled with bouts of proboscis extension sleep. Notably, experimental sealing of the anus in *ctrl* flies phenocopied the behavioural and physiological impairments observed in *ap^ΔLSE^* mutants, reinforcing the necessity of meconium excretion for normal feeding initiation. This study offers new insights into the conserved hierarchical events regulating juvenile fly behaviour, and highlights a critical physiological process necessary for the onset of adult feeding.

### Resolving a 110-year-old enigma: *ap* and the precocious adult death phenotype

In 1914, Metz reported on the first *ap* allele (*ap^1^*). It was characterized by pleiotropic phenotypes such as the absence of wings and halteres, premature adult death and sterility (Metz, 1914). While the role of *ap* in wing development has been extensively studied (reviewed in Irvine and Rauskolb 2001), the mechanisms underlying the precocious adult death phenotype remained unclear. In this study, we identified a *cis*-regulatory enhancer, the lifespan enhancer, or LSE, which is critical for adult survival by ensuring proper rectal papillae formation. In its absence, papilla precursors fail to undergo proper development, leading to the formation of the Reinger’s knot, a hindgut obstruction that ultimately prevents meconium excretion, disrupts adult feeding and promotes a proboscis extension sleep state.

Our findings suggest that *ap* plays a critical role in the morphogenesis of rectal papillae during early pupal development. Notably, the timing of papillae formation coincides with two rounds of mitotic division in the polyploid papilla precursor cells (Fig. 4; Fox et al., 2010; Schoenfelder et al., 2014). In *ap^ΔLSE^*flies, these precursor cells remain clustered together within the Reinger’s knot. Given that the LSE drives *ap* expression in only two of the four rectal papillae, we hypothesize that *ap* regulates the expression of cell surface proteins required for the separation of the papilla precursors into four papillae. This might be similar to the role of *ap* in wing compartmentalization, where it contributes to the separation of dorsal and ventral wing cells (Blair et al., 1994). While Notch signaling is also known to be essential for papillae formation (Fox et al. 2010; Schoenfelder et al. 2014), the distinct phenotypes observed in Notch-RNAi and *ap^ΔLSE^*flies suggest that they play distinct roles. Future studies shall investigate the specific molecular interactions between *ap* and *Notch* signaling in the context of rectal papillae development.

Interestingly, we also found that female sterility, which is observed in all *ap^null^* alleles, is a secondary effect of the conditions induced by the Reinger’s knot in *ap^ΔLSE^* flies (Reinger et al, in preparation; King and Sang 1958; Butterworth and King 1964, 1965; King and Bodenstein 1965; Butterworth 1972; Postlethwait and Weiser 1973; Richard et al 1993; Aguila et al 2007). Therefore, it seems that under stressful metabolic conditions, flies make critical metabolic decisions on whether to bear the cost of reproduction at the possible expense of survival (Meiselman et al., 2018).

### A *Drosophila* model for ileus disease: parallels with human gastrointestinal dysfunction

In medicine, ileus is defined as a temporary cessation of intestinal peristalsis (Plusczyk et al. 2006). In practice, a distinction is made between mechanical and paralytic ileus. A paralytic ileus can have various pathological causes that lead to an impairment of intestinal peristalsis. In contrast, intestinal peristalsis is not impaired in mechanical ileus, at least initially. Rather, it is the result of a mechanical obstacle to passage. Regardless of its etiology, a first pathophysiological indication of a mechanical ileus is a palpable dilatation of the intestinal wall due to a several fold increase in intraluminal pressure. A second important diagnostic feature of mechanical ileus is hyperperistalsis, by which the intestine attempts to push its contents past the intestinal obstruction. Within the first 24 hours, the mechanical stress leads to a considerable reduction in water and salt ion absorption from the intestinal lumen. In addition, the reduced blood circulation leads to hypoxia and paralysis of the energy-consuming molecular transport systems. A vicious circle is set in motion that further increases the intraluminal pressure. The intestinal epithelium becomes permeable, leading to bleeding, peritonitis and translocation of intestinal bacteria. An untreated ileus inevitably leads to perforation of the intestinal wall with fatal and life-threatening consequences for the patient (Ansari, 2024; Wang et al., 2024).

Our detailed analysis of the phenotypes detected in *ap^ΔLSE^* flies suggests that there are some striking parallels to humans suffering from a mechanical ileus. In both, reduced appetite, absolute constipation and intestinal bloating are first indicators for the pathophysiological condition. Another important diagnostic feature for a mechanical ileus is hyperperistalsis, a compensatory mechanism that attempts to force intestinal contents past the obstruction, eventually leading to increased intraluminal pressure, and, if unresolved, tissue rupture with fatal consequences for the patient (Ansari, 2024; Wang et al., 2024). A similar phenomenon may be occurring in *ap^ΔLSE^* flies, where the sleep state known as PES (van Alphen et al., 2021) could not only minimize energy consumption and increase haemolymph circulation but also stimulate gut peristalsis in an attempt to expel the obstructing meconium.

These findings establish *Drosophila* as a genetically tractable model for investigating ileus-like conditions and physiological aspects of gastrointestinal dysfunction.

## Conclusion

Our study not only resolves a century-old mystery surrounding the adult precocious death phenotype in *ap* mutants, but also provides fundamental insights between the coupling of meconium excretion, feeding initiation and PES in *Drosophila melanogaster*. Moreover, the discovery of a genetic mechanism underlying meconium-induced constipation in *ap^ΔLSE^*mutants establishes a new model for studying gastrointestinal obstruction and its physiological consequences in *Drosophila*. However, the mechanisms linking meconium retention to feeding inhibition remain unresolved. One possibility is that the presence of the meconium in the gut provides physiological or mechanical signals indicating satiety, thereby delaying the drive to ingest food. This process will likely involve interactions between the digestive, endocrine and nervous systems, to coordinate the onset of feeding when needed. Future studies will be needed to establish the mechanisms that couple these processes.

## Methods

### Drosophila strains and culture

Flies were housed in a 25°C incubator with 60% humidity under a 12h light:12h dark cycle, and reared on a standard cornmeal agar diet composed of 0.07% w/v granulated agar-agar, 0.1% w/v corn meal, 0.06% w/v granulated sugar, 0.015% w/v granulated dry yeast, 0.003% w/v granulated Nipagin. The solidified medium was supplemented with a drop of fresh baker’s yeast unless otherwise stated.

*ap^DG1^,* also referred to as *ap^DG^*, was described in Gohl et al. (2008). *ap^c1.4b^*, *ap^R1^*(also called *ap^attPΔEnh^*) and *ap^DG3^* are described in Bieli et al 2015a. *ap^R2^* is described in Aguilar et al. (2023). *P{UAS-mCherry^NLS^}* is described in Caussinus et al (2008). *UAS-ci^Rep^* (*P{UAS-ci^76^}*) was obtained from Pascal Thérond (Institut de Biologie Valrose; Aza-Blanc et al., 1997). *byn^Gal4^* was shared by Donald T. Fox (Duke University; Iwaki and Lengyel, 2002), *y M{vas-int.Dm}zh-2A w* by Johannes Bischof (University of Zürich; Bischof et al., 2007), and *P{UAS-ap}* by Marco Milán (IRB Barcelona; Milán and Cohen, 1998).

The following stocks were obtained from the Bloomington Drosophila stock center: *G-TRACE* (#28281), *ap^md544^* (#3041), *P{tub-Gal80^ts^}* (#7019), *P{UAS-mCD8::GFP}* (#32186), *Dl^Gal^* (#67047) and *M{vas-Cas9}ZH-2A* (#51323). *PBac{RB}e01573* was purchased from Exelixis (Harvard; no longer available).

#### Construction of byn^Gal4^ UAS-mCherry^NLS^

*byn^Gal4^*was recombined with *P{UAS-mCherry^NLS^}*.

#### Construction of *byn^Gal4^* tubGal80^ts^

*byn^Gal4^*was recombined with *P{tub-Gal80^ts^}*.

#### Construction of *byn^Gal4^UAS-ci^Rep^*

*byn^Gal4^*was recombined with *P{UAS-ci^76^}*.

#### Stocks generated for temperature shifts experiments

*y w; P{tub-Gal80^ts^} ap^md544^/ CyO, Dfd::YFP. ap^md544^* (Calleja et al., 1996) is a strong *ap* hypomorph as well as a *bona fide ap^Gal4^*driver. *ap^md544^ / ap^DG3^* animals have almost no wings and halteres and die precociously.

*y w; P{UAS-mCD8::GFP} ap^DG3^ P{UAS-ap} / CyO, Dfd::YFP*.

*y w; P{UAS-mCD8::GFP} ap^DG3^ / CyO, Dfd::YFP* was used as control.

### New *ap* alleles established for this study

#### Generation of the *ap^MS1^* landing site

Preliminary observations suggested that a regulatory region required for adult survival is likely located in or between apCON1 and the proximal side of the apE wing enhancer (M. M., unpublished). Therefore, this region was deleted and replaced with a landing site for the ɸC31 integrase system and an FRT using CRISPR/Cas9-mediated homology directed repair (Gratz et al., 2013). In preparation for this approach, two plasmids containing one gRNA each were constructed in the following way. Plasmid pU6-BbsI-chiRNA (Gratz et al., 2013) was cut with the restriction enzyme BbsI and ligated with the following oligonucleotides:

gRNA “CRISPR L” located in apCON1:

5’-CTTCGTAAACTCAAGTATCGGAAT

3’-ATTTGAGTTCATAGCCTTACAAA-5’

gRNA “CRISPR R” located in apCON2:

5’-CTTCGAAGCTGCACTATGACTGAG-3’

3’-CTTCGACGTGATACTGACTCCAAA-5’

The newly generated plasmids are pMS393 (for “CRISPR L”) and pMS394 (for “CRISPR R”). Donor plasmid pMS398 was designed *in silico* and synthesized by Genewiz (Azenta Life Sciences). It consists of two ∼700 bp homology arms, which flank a cassette consisting of [55 bp attP]-[34 bp FRT]-[517 bp Dad13 enhancer]-[34 bp FRT]. The wing-specific Dad13 enhancer serves as selectable marker as described in Aguilar et al. (2023).

Donor plasmid pMS398 (at 500 ng/l) was injected together with the two gRNA producing plasmids (pMS393 and pMS394; at 100 ng/l) into embryos with genotype *y^1^ M{vas-Cas9}zh-2A w^1118^; +*. Successful HDR events lead to the deletion of 3143 bp from the *ap* locus and insertion of the “dominant marker” Dad13. Adult injectees were crossed with *y w* flies and their progeny was screened for the weak dominant wing phenotype generated by the presence of the Dad13 enhancer in the *ap* locus (Aguilar et al., 2023). Candidates were verified by PCR and sequencing and a *y w; ap^MS1+Dad13^/ CyO, Dfd::YFP* stock was established. In a second step, the Dad13 enhancer was removed by Flipase-mediated recombination (Golic and Golic, 1996). The new allele is called *ap^MS1^*(not shown). Homozygous flies show the precocious adult death phenotype but have normal wings. The sequences on the proximal and distal sides of this deletion are:

proximal side (attP underlined):

TCCCGAACGAAAAGAAAAGA<u>GGAGTAGTGCCCCAACTGGGGTAACCTTTGAGTTCTCTCAGTTGGGGGCGTAGGC</u>

distal side (FRT underlined):

<u>GAAGTTCCTATTCTCTAGAAAGTATAGGAACTTC</u>CGTAGTGTGCAATGCACACT

#### Generation of the *ap^MS2^* landing site (*ap^ΔLSE^*; Supp. Figure 1D)

Although *ap^MS1^* flies are short-lived, complementation data suggested that not all of the life span enhancer is missing on this chromosome (M. M., unpublished). Therefore, the *ap^MS1^*deletion was extended on its distal side. This was achieved by Flipase-induced recombination between the FRTs present on *ap^MS1^* and *ap^R2^* (Supp. Figure 1F). This leads to a deletion of 3794 bp. The allele is called *ap^MS2^*. Please note that it is referred to as *ap^ΔLSE^* in the main text and in all figures. The sequence on the distal side of this deletion is (proximal side same as above):

Distal side (FRT underlined):

<u>GAAGTTCCTATTCTCTAGAAAGTATAGGAACTTCT</u>ATAATTTGACCGCAATTTT

As *ap^MS1^*, homozygous *ap^MS2^* flies show the precocious adult death phenotype but they also lack wings and halteres because the apE wing enhancer is deleted as well (Bieli et al., 2015a, b).

#### Construction of *ap^MS2+minLSE^* (*ap^minLSE^*; Supp. Figure 1G)

The LSE (562 bp fragment) was cloned in reentry vector DB345 (Bieli et al., 2015b) and plasmid CR24 was obtained. It was injected into *y w vas-integrase; ap^MS2^ / CyO* embryos. Successful integration into the *ap^MS2^* landing site was detected by the expression of the *yellow* reporter. The *y^+^* marker was subsequently removed by Flp-treatment (Golic and Golic, 1996). Flip-out candidates were verified by PCR and sequencing and a balanced stock was established.

#### Construction of *ap^c1.4b-Gal4^* (*ap^Gal4^*; Supp. Figure 1H)

*ap^c1.4b-Gal4^* was obtained by injection of plasmid pMS399 into *y w vas-integrase; ap^c1.4b^ / CyO* embryos (Supp. Fig. 1A). pMS399 contains a *mini-yellow* marker devoid of all *yellow* enhancers. Injectees were crossed with *y w* partners. Transgenic flies could be detected by their y^+^ wings. A *y w; ap^c1.4b-Gal4^ / CyO* stock was established and crucial parts of the insert verified by sequencing. Note that reporter gene expression patterns generated by the classic *ap^md544^* allele (Calleja et al., 1996) and *ap^c1.4b-Gal4^* are indistinguishable. But unlike *ap^md544^*, hemizygous *ap^c1.4b-Gal4^* flies are well viable and fertile and have normal wings and halteres. Please note that it is referred to as *ap^Gal4^*in the main text and in all figures.

The Gal4 gene in this and the following two *ap^Gal4^* alleles is under the control of a synthetic core promoter. It consists of a modified 155 bp fragment originating from the *evenskipped* gene and contains consensus motifs known to be present in potent Drosophila promoters. They are the TATA-box, the initiator element, the motive ten element and the downstream promoter element (Pfeiffer et al., 2008).

#### Construction of *ap^DG1-Gal4^* (*ap^ΔLSE-Gal4^*; Supp. Figure 1I)

*ap^DG1-Gal4^* was generated by Flp-mediated recombination between FRT sites present in *ap^c1.4b-Gal4^* and *PBac{RB}e01573* (Thibault et al., 2004). The transgene insert *PBac{RB}e01573* marks the distal end point of the 27 kb intergenic spacer. The same strategy was previously used to generate *ap^DG1^* (Supp. Fig. 1C; Gohl et al., 2008). Hence, all *ap* enhancers located in the intergenic spacer including the LSE are lost. With respect to survival, *ap^DG1-Gal4^*behaves like *ap^ΔLSE^*. Note that in *ap^DG1-Gal4^*, the *mini-yellow* marker of *ap^c1.4b-Gal4^* and the *mini-white* marker of *PBac{RB}e01573* have been deleted by the Flp-procedure. Also note that it is referred to as *ap^ΔLSE-Gal4^* in the main text and in all figures.

#### Construction of *ap^R1+LSE-Gal4^* (*ap^LSE-Gal4^*; Supp. Figure 1J)

This Gal4 driver was obtained by insertion of plasmid pCR38 into the *ap^R1^* landing site (Supp. Fig. 1E). In *ap^R1+LSE-Gal4^*, the whole 27 kb intergenic spacer is replaced by the 562 bp minimal LSE and the Gal4 gene. *ap^R1+LSE-Gal4^* flies are viable and fertile, indicating that the LSE is complementing for the deleted intergenic spacer. This suggests that that the LSE also activates the Gal4 gene accordingly. Note that the *yellow* marker present on pCR38 remains in place and is located between the 3′end of Gal4 and the 5′end of *l(2)09851*. This yellow marker contains the body color enhancer but not the wing enhancer of the *yellow* gene. Note that *ap^R1+LSE-Gal4^* is referred to as *ap^LSE-Gal4^*in the main text and in all figures.

### Sample preparation and immunohistochemistry

#### G-TRACE

In order to follow *ap* expression throughout different stages of development, Gal4 driver lines were crossed with the real-time and clonal expression stock (G-TRACE; Evans et al., 2009). Dechorionated embryos or dissected tissue samples were fixed with 4% paraformaldehyde in PBS for 25 min at room temperature (RT). Samples were washed 3x for 20 min each with PBS at RT and finally mounted in Vectashield containing DAPI (Vector Laboratories). Confocal imaging was performed using a Leica SP5 microscope or an Olympus SpinD. Image processing was done with the ImageJ software.

#### Immunostaining

In order to verify the specificity of the ap expression generated by G-TRACE stainings, antibody stainings against Ap was performed. Standard immune-detection protocols were followed to image late stage embryos. Primary antibody used in this study was rabbit α-Ap (1:800-1000, Bieli et al., 2015b). Secondary antibody was α-rabbit Alexa Fluor 488 used at 1:500. Samples were mounted in Vectashield containing DAPI. Confocal imaging was performed using a Leica SP5 microscope or an Olympus SpinD. Image processing was done with the Omero or with the ImageJ software.

#### Ampulla preparations

Adult ampullae were dissected in 1x PBS on a siliconized slide and mounted in 1x PBS / 50% Glycerol. In order to prevent crushing of the tissue samples, coverslips with tiny playdough feet were used. Preparations were immediately inspected with a Leica DM2700M microscope equipped with Normarski optics and imaged with a Leica Flexacam C3 camera.

### micro-computed tomography (µ-CT)

Micro-computed tomography was used as a non-invasive imaging method to achieve high-resolution visualization of the pupal and adult hindgut. μ-CT scans were done as previously described (Schoborg et al., 2019; Blackie et al., 2024). In short, female pupae were staged and pooled in Eppendorf tubes containing 1 ml of 0.5% TritonX-100 (Sigma, T8787) in PBS, incubated at 100°C for 20 sec in a heatblock, and then allowed to cool down at RT for 5 min. This kills the pupae immediately so that they do not progress to later developmental stages and partially permeabilizes the cuticle. One-day-old female virgins were anesthetized with CO_2_. 5 - 50 flies were pooled in a 1.5 ml Eppendorf tube. 1 ml of 0.5% TritonX-100 in PBS (PBST) was then added for around 5 min or until flies sunk to the bottom of the tube. For both stages, PBST was exchanged for 1 ml of Bouin′s solution (Sigma-Aldrich, HT10132) for 16-24 hours to fix the samples, which were then washed 3x for 30 min and then again overnight with PBST. They were stained in 1 ml of 1:1 Lugol′s solution (Sigma, 62650) in ultrapure H_2_O for a minimum of 3 days. To remove the staining solution, the samples were washed with ultrapure water once and then stored at RT in ultrapure water until ready to scan. Samples were finally mounted head down in a 10μl pipette tip filled with ultrapure water and the ends were sealed with parafilm. 5 - 10 flies were mounted per tip.

Pupae and flies were imaged on a Bruker Skyscan 1272 with the following settings: 40 kV, 110 μA, 4 W, CMOS camera scanning at a 2.95 μm pixel size, 0.3–0.35 rotation step, 30 μm random movement and four frame averaging. Images were reconstructed using the Bruker NRecon software, then images were further processed by background subtraction and Gaussian smoothing in FIJI v.2.14.0/1.54f.

The software ITK-snap v3.8.0 was used to manually segment each intestine with the polygon tool. Organ perimeter were segmented every 15-20 slices in the axial plane and the morphological interpolation tool was used to fill in the spaces. Smoothing factors were added when needed. Manual corrections were performed using the adaptive paintbrush tool. For 3D visualization, segmentations were converted into meshes and were further processed with the software Meshlab. Meshes were processed with the quadric edge collapse decimation to reduce the number of faces to 10% and/or smoothened using HC Laplacian smoothing.

### Behaviour Assays

#### Survival Assays

##### Quantification of survival rates

Unless otherwise stated, survival rates for the various genotypes were carefully scored over a period of 18 days. Flies of both sexes were collected during one day and kept at 25°C in pools of 20-30 flies. Flies were kept in glass vials (∼56 cm^3^) on a normal cornmeal agar diet not supplemented with fresh baker’s yeast. Flies were transferred to fresh tubes at least every second day without CO_2_ anesthesia. The number of surviving flies was scored every third day. Survival curves are plotted as survival in % ([survivors on day X]/[total of flies scored]x100) versus day X after eclosion.

##### Survival under high salt conditions

As above but standard food was supplemented to reach a final concentration of 250 mM NaCl.

##### Survival of starved *y w* flies

Freshly eclosed *y w* females and males were collected separately and directly transferred to starvation tubes (1.5% agar in water) or vials containing standard food. Flies kept on normal food were aged for 5 days before they were transferred to starvation tubes. Survival was scored until the last fly died.

##### Survival of butt-plugged *y w* flies

Butt-plugged flies were produced by sealing the anus of freshly eclosed *y w* females and males with a drop of super glue (Pattex, #2804584) under CO_2_ anesthesia. Thereafter, females and males were kept in separate tubes containing standard diet. Survival was scored until the last fly died. During this period, flies were transferred into fresh tubes at the latest every second day.

#### Feeding, excretion and proboscis extension assays

##### flyPAD

In order to test if the *ap* mutant flies are interacting with food or not, flyPAD assays were performed (Itskov et al., 2014; Hadjieconomou et al., 2020). Food pellet composition was: 5% w/v sucrose (granulated sugar, Tate & Lyle), 10% w/v Brewer’s yeast (#903312, MP Biomedicals), 1.5% w/v agar (Sigma A7002) supplemented with Nipagin (Sigma H5501; 30 mL/L of 10% w/v Nipagin in 95% EtOH). Nipagin was added once the food had cooled down to < 60°C. The food was poured in 5 cm petri dishes. Solid food pellets were punched out using a P1000 cut tip to match the exact diameter of the central electrode patch. One-day-old virgins were then individually transferred to the flyPAD arena by mouth aspiration. Half of the wells were filled with a food pellet, the other one was left empty. Virgins were allowed to feed for 1.5 hours at 25 °C. Only the last 60 min were used for quantification. flyPAD assays were only carried out during the feeding peak between 10 am to 2 pm. The total number of sips per fly was acquired with the Bonsai framework and further analyzed in MATLAB.

##### Blue food squash assay

The amount of ingested food was quantified with food containing 1% FCF blue (Hadjieconomou et al., 2020). In short, virgins were collected over one day. Later in the afternoon, they were transferred to starvation tubes (1.5% agar in water). On the following day, after around 16 hours of starvation, the flies were transferred into tubes containing blue food and allowed to feed for a maximum of 20 min. They were subsequently frozen in liquid nitrogen and transferred in groups of three to a clean 2 ml PCR tube (Eppendorf, 22431048) with 0.5 ml of water and a 5 mm stainless-steel metal bead (QIAGEN, 69989). The flies were then homogenized using a QIAGEN TissueLyser II for 90 s at 30 Hz. The samples were centrifuged at 10,000g for 5 – 10 min. 200μL of the supernatant per sample was directly transferred into individual wells of a 96-well, flat bottom, optically clear plate (Thermo Fisher Sterilin, 611F96). A BMG Labtech FLUOstar Omega plate reader was used to measure dye content by reading the absorbance at 629 nm. We used a standard curve of pure FCF blue dye to calculate the dye content ingested per fly.

##### Crop-mount assay

Flies were prepared the same way as described for the blue food squash assay. Instead of homogenizing them, their foregut and crop were quickly dissected and mounted on a poly-lysine-coated slide. With a normal bright field microscope (Leica DM2700M) attached to a camera (Leica Flexacam C3), the crops were imaged. Further, their area was manually analyzed by using the ImageJ software.

##### FlyPoo Assay

Virgins were prepared the same way as described for the blue food squash assay. Instead of putting them into vials containing blue food, 30 flies were transferred into individual petri dishes (Sarstedt, #82.1194.500) containing a wedge of blue diet. They were allowed to eat and poop for 2 hours. Afterwards, the flies were removed and the excreted feces on the petri dishes were manually counted.

##### Meconium Excretion Assay

To characterise the rate and dynamics of meconium excretion, the following assay was performed: freshly eclosed females and males (max. 15 minutes old) were collected with the shortest possible CO_2_ anesthesia. Between 20 - 30 flies for each sex were transferred into petri dishes (Sarstedt, #82.1194.500) containing a wedge of standard food or in an empty petri dish without any food. For each time point, another batch of flies was used. The total number of excreted feces was manually counted for each sex and petri dish. The total number of excreta was then divided by the number of flies present in each petri dish.

##### Adult initial feeding assay

To analyze when juvenile flies start to eat after eclosion, the adult initial feeding assay was performed. Flies were prepared the same way as for the meconium excretion assay. Instead of petri dishes supplemented with standard food, 10 - 20 flies were transferred into a petri dish (Sarstedt, #82.1194.500) containing a wedge of blue diet (same composition as for the blue food squash assay). Every hour, the first appearance of blue food in the flýs intestine was scored by eye. For the time points 10 and 24 hours after eclosion, new batches of flies were used. The number of excreted meconium as well as the number of excreted blue diet was manually counted.

##### Proboscis extension assays

For video-assisted analysis of the various genotypes, virgins were collected over one day and kept in vials filled with the same food as described in the flyPAD section until their behaviour was recorded on the following morning from 9 am to 10 am or 10 am to 11 am (4 individual flies per 1 hr interval). During recording, flies have unrestricted access to normal food (for food content see flyPAD assay). For experiments described in Fig. 2J-K, starved and well fed *ctrl* flies were put on starvation food (1.5 % agar). The behaviour experiments were done using a custom-build 3D-printed chamber comprising 4 individual small chambers as described previously (Cury and Axel, 2023). In brief, flies were placed in the chamber and recorded for a period of one hour with an aCA2440-75um camera (Basler) mounted with a 0.5X 2/3” Telecentric lens (GoldTL ^TM^, Edmund optic) equipped with a near infrared bandpass filter (Midopt BP850-58). Illumination was achieved by directing the light from the infrared LED (M850L3, Thorlabs) onto a glass surface positioned at a 45° angle above the chamber, enabling reflection of the light downwards. The flies were recorded at a resolution of 142 px/mm (1386 × 1066 pixels). Manual annotation of the proboscis extension (PE) was conducted by using Behaviour Observation Research Interactive Software (BORIS, Friard and Gamba, 2016). Three types of PE were identified: PE on food, PE while moving and PE while immobile. The start of a PE event was considered as the first frame where labellum extension was observed, and the end was defined as the frame after the proboscis was completely retracted. For the immobility analysis, videos were analyzed using SLEAP (Pereira et al., 2022) to track the head, thorax and abdomen tip. The fly was considered immobile when there was no movement of all three points. This method excludes periods of grooming, as either the head or the abdomen moves during this behaviour. A fly is considered to enter a sleep period if it remains immobile for more than 5 min. Data was analyzed using an in-house Python script developed for this study.

#### Temperature shifts experiments

##### Determining the temporal requirement for *ap* expression with respect to the precocious adult death phenotype

To address the temporal requirement for *ap* expression with respect to the adult precocious death phenotype, the following experimental procedure was followed. *y w; tub-Gal80^ts^ ap^md544^ / CyO, Dfd::YFP* virgins were crossed with *y w; UAS-mCD8::GFP ap^DG3^ UAS-ap / CyO, Dfd::YFP* males. Embryos were collected in a fresh culture tube containing standard cornmeal agar supplemented with fresh baker’s yeast for 24 hours at 18°C. Parents were removed and the collected animals were further aged at 18°C for 96, 120, 144, 168, 192, 216, 240, 264, 288, 312 or 336 hours before cultures were shifted to 29°C and kept at this temperature till adulthood. Then, the survival of emerging adults with the genotype *UAS-CD8::GFP ap^DG3^ UAS-ap* / *ap^md544^* tub^Gal80ts^ was carefully scored.

*UAS-mCD8::GFP ap^DG3^ UAS-ap* / *ap^md544^ tub-Gal80^ts^* animals continuously grown at 18°C did not show a GFP pattern in 3^rd^ instar larvae. Furthermore, adults did not form wings and showed the precious adult death phenotype. At 29°C constantly, an ap-like GFP pattern was seen in 3^rd^ instar larvae, indicating that Gal80^ts^ is no longer functional and that Gal4 activates the UAS-transgenes. Consequently, adults developed normal wings and survived well (Supp. Table 3). A control experiment in the absence of *UAS-ap* failed to rescue *ap^DG3^* / *ap^md544^* phenotypes at 29°C (not shown).

##### Determining the temporal requirement for the *byn^Gal4^>ci^Rep^*effect on papillae development

*byn^Gal4^ tub-Gal80^ts^ / TM6C* virgins were crossed with homozygous *y w; UAS-ci^Rep^* males. When *byn^Gal4^ tub-Gal80^ts^ / UAS-ci^Rep^* animals were constantly grown at 18°C, adults survived well. Dissected female ampullae contained four papillae. When constantly grown at 29°C, adults also survived well but they had no papillae (Supp. Table 4). For the temperature shifts, the experimental procedure was as follows. The same cross as above was set up. Embryos were collected in a fresh culture tube containing standard cornmeal agar supplemented with fresh baker’s yeast for 24 hours at 18°C. Parents were removed and the collected animals aged at 18°C for 48, 96, 144, 192, 240, 264 or 288 hours before cultures were shifted to 29°C. Cultures were then kept at this temperature till adulthood. Once adults started to emerge, the phenotypic classes were scored (Sb vs. non-Sb flies). The two classes emerged at the expected Mendelian ratio (1:1; not shown). In addition, the survival and fertility of non-Sb flies (*byn^Gal4^ tub-Gal80^ts^ / UAS-ci^Rep^*) was scored, and the hindguts of several females were inspected for the presence or absence of rectal papillae.

## Acknowledgments

C.R. would like to thank Ludovico Maggi, Gustavo Aguilar and Sophie Schnider for fruitful discussions and input of new ideas bringing this project forward. Thanks are also due to David Ruel for the quantification of the sleep videos. A special thank you goes to Paola Cognigni and Gerrit Linneweber for their idea about sealing the anus of adult flies. We are indebted to Dragana Rogulja for insightful comments and discussions about the PES phenotype. We thank Dr. med. Andres Heigl (Kantonsspital Baselland) who clarified our view on human ileus. The Imaging Core Facility of the Biozentrum Basel is also acknowledged. We would also like to thank Pascal Thérond, Donald T. Fox, Johannes Bischof, Marco Milán and the Bloomington Stock Center for sending flies. The Biozentrum fly community is grateful to Gina Evora, Consuelo Zuluaga Gomez, Adela Garzia and Patrick Groelly for constant and reliable supply of world’s best fly food. The work in the laboratory of M.A. was supported by grants from the Swiss National Science Foundation (310030_192659 and 310030B_176400) and by funds from the Kanton Basel-Stadt and Basel-Land. The work in the laboratory of I.M-A. was funded by an ERC Advanced Grant (ERCAdG 787470 ‘IntraGutSex’), a UKRI Horizon Europe guarantee grant (EPSRC-ERC EP/Y03298/1), MRC intramural funding and the Francis Crick Institute, which receives its core funding from Cancer Research UK (FC001317, FC001175), the UK Medical Research Council (FC001317, FC001175) and the Wellcome Trust (FC001317, FC001175). The work in the laboratory of A.K. was supported by the Swiss State Secretariat for Education, Research and Innovation (SERI) (M822.00025, SERI-funded ERC Starting Grant) and by funds from the Kanton Basel-Stadt and Basel-Land.

## Author contribution

C.R. conducted all experiments unless otherwise stated. M.M. conducted the dissection and brightfield imaging of all ampulla and established all fly lines. C.R. and M.M. conducted the temperature shift experiments and *ap* survival assays. M.S. designed and generated the *ap^MS2^*landing site as well as *ap^Gal4^*. A.M., H.G. and C.R. conducted high resolution proboscis extension video assays. A.M. and H.G. did the quantification of the data and provided figures. P.G., D.H. and C.R. conducted the feeding related experiments, P.G. provided figures. P.G. did the muscle stainings and provided figures. L.B. and C.R. conducted the microCT scans, L.B. did the analysis of the scans, the segmentations and provided figures. D.H. conducted the stainings of larval and adult guts and provided figures. M.A. acquiring funding for the work. C.R., M.M. and A.K. wrote the original draft. All other authors read, edited, commented on and approved the final manuscript.

## References

Aguilar, G., Sickmann, M., Bieli, D., Born, G., Affolter, M., Müller, M., 2023. Transcriptional control of compartmental boundary positioning during Drosophila wing development. bioRxiv 2023.08.05.552106. 10.1101/2023.08.05.552106

Aguila, J.R., Suszko, J., Gibbs, A.G., Hoshizaki, D.K., 2007. The role of larval fat cells in adult Drosophila melanogaster. J. Exp. Biol. 210, 956–963. 10.1242/jeb.001586

Ahanya, S.N., Lakshmanan, J., Morgan, B.L.G., Ross, M.G., 2005. Meconium passage in utero: mechanisms, consequences, and management. Obstet. Gynecol. Surv. 60, 45–56; quiz 73–74. 10.1097/01.ogx.0000149659.89530.c2

Alcantara, I.C., Tapia, A.P.M., Aponte, Y., Krashes, M.J., 2022. Acts of appetite: neural circuits governing the appetitive, consummatory, and terminating phases of feeding. Nat. Metab. 4, 836–847. 10.1038/s42255-022-00611-y

Apidianakis and Rahme, 2011. *Drosophila melanogaster* as a model for human intestinal infection and pathology. Dis Model Mech (2011) 4 (1): 21–30. 10.1242/dmm.003970

Ansari, P., 2024. Ileus. https://www.msdmanuals.com/professional/gastrointestinal-disorders/acute-abdomen-and-surgical-gastroenterology/ileus

Aza-Blanc, P., Ramírez-Weber, F.A., Laget, M.P., Schwartz, C., Kornberg, T.B., 1997. Proteolysis that is inhibited by hedgehog targets Cubitus interruptus protein to the nucleus and converts it to a repressor. Cell 89, 1043–1053. 10.1016/s0092-8674(00)80292-5

Bateman et al 2006. Site-specific transformation of Drosophila via phiC31 integrase-mediated cassette exchange. Genetics 173(2):769–77. doi: 10.1534/genetics.106.056945

Bergman, C.M., Pfeiffer, B.D., Rincón-Limas, D.E., Hoskins, R.A., Gnirke, A., Mungall, C.J., Wang, A.M., Kronmiller, B., Pacleb, J., Park, S., Stapleton, M., Wan, K., George, R.A., de Jong, P.J., Botas, J., Rubin, G.M., Celniker, S.E., 2002. Assessing the impact of comparative genomic sequence data on the functional annotation of the Drosophila genome. Genome Biol. 3, RESEARCH0086. 10.1186/gb-2002-3-12-research0086

Betley, J.N., Xu, S., Cao, Z.F.H., Gong, R., Magnus, C.J., Yu, Y., Sternson, S.M., 2015. Neurons for hunger and thirst transmit a negative-valence teaching signal. Nature 521, 180–185. 10.1038/nature14416

Bieli, D., Kanca, O., Gohl, D., Denes, A., Schedl, P., Affolter, M., Müller, M., 2015a. The Drosophila melanogaster Mutants apblot and apXasta Affect an Essential apterous Wing Enhancer. G3 Bethesda Md 5, 1129–1143. 10.1534/g3.115.017707

Bieli, D., Kanca, O., Requena, D., Hamaratoglu, F., Gohl, D., Schedl, P., Affolter, M., Slattery, M., Müller, M., Estella, C., 2015b. Establishment of a Developmental Compartment Requires Interactions between Three Synergistic Cis-regulatory Modules. PLOS Genet. 11, e1005376. 10.1371/journal.pgen.1005376

Bischof, J., Maeda, R.K., Hediger, M., Karch, F., Basler, K., 2007. An optimized transgenesis system for Drosophila using germ-line-specific φC31 integrases. Proc. Natl. Acad. Sci. 104, 3312–3317. 10.1073/pnas.0611511104

Blackie, L., Gaspar, P., Mosleh, S., Lushchak, O., Kong, L., Jin, Y., Zielinska, A.P., Cao, B., Mineo, A., Silva, B., Ameku, T., Lim, S.E., Mao, Y., Prieto-Godino, L., Schoborg, T., Varela, M., Mahadevan, L., Miguel-Aliaga, I., 2024. The sex of organ geometry. Nature 630, 392–400. 10.1038/s41586-024-07463-4

Blair, S.S., Brower, D.L., Thomas, J.B., Zavortink, M., 1994. The role of apterous in the control of dorsoventral compartmentalization and PS integrin gene expression in the developing wing of Drosophila. Dev. Camb. Engl. 120, 1805–1815. 10.1242/dev.120.7.1805

Bownes, M., 1982. Hormonal and genetic regulation of vitellogenesis in Drosophila. Q. Rev. Biol. 57, 247–274. 10.1086/412802

Buchon, N., Broderick, N.A., Lemaitre, B., 2013. Gut homeostasis in a microbial world: insights from Drosophila melanogaster. Nat. Rev. Microbiol. 11, 615–626. 10.1038/nrmicro3074

Butterworth and King 1964. The fat body of wild-type and of various mutant apterous adults. DIS 39:82.

Butterworth and King 1965. The developmental genetics of apterous mutants of Drosophila melanogaster. Genetics 52, 1153–1174. 10.1093/genetics/52.6.1153

Butterworth F. M. (1972). Adipose tissue of Drosophila melanogaster. V. Genetic and experimental studies of an extrinsic influence on the rate of cell death in the larval fat body. Developmental biology, 28(2), 311–325. 10.1016/0012-1606(72)90016-4

Calleja, M., Moreno, E., Pelaz, S., Morata, G., 1996. Visualization of Gene Expression in Living Adult Drosophila. Science 274, 252–255. 10.1126/science.274.5285.252

Capovilla, M., Kambris, Z., Botas, J., 2001. Direct regulation of the muscle-identity gene apterous by a Hox protein in the somatic mesoderm. Dev. Camb. Engl. 128, 1221–1230. 10.1242/dev.128.8.1221

Caussinus et al 2008. Tip-cell migration controls stalk-cell intercalation during Drosophila tracheal tube elongation. Current biology, 18(22), 1727–34. 10.1016/j.cub.2008.10.062.

Chen, Y., Lin, Y.-C., Kuo, T.-W., Knight, Z.A., 2015. Sensory Detection of Food Rapidly Modulates Arcuate Feeding Circuits. Cell 160, 829–841. 10.1016/j.cell.2015.01.033

Cohen, B., McGuffin, M.E., Pfeifle, C., Segal, D., Cohen, S.M., 1992. apterous, a gene required for imaginal disc development in Drosophila encodes a member of the LIM family of developmental regulatory proteins. Genes Dev. 6, 715–729. 10.1101/gad.6.5.715

Cohen, E., Sawyer, J.K., Peterson, N.G., Dow, J.A.T., Fox, D.T., 2020. Physiology, Development, and Disease Modeling in the Drosophila Excretory System. Genetics 214, 235–264. 10.1534/genetics.119.302289

Cury and Axel, 2023. Flexible neural control of transition points within the egg-laying behavioral sequence in *Drosophila*. Nature Neuroscience | Volume 26, 1054–1067. 10.1038/s41593-023-01332-5

de Taffin, M., Carrier, Y., Dubois, L., Bataillé, L., Painset, A., Le Gras, S., Jost, B., Crozatier, M., Vincent, A., 2015. Genome-Wide Mapping of Collier In Vivo Binding Sites Highlights Its Hierarchical Position in Different Transcription Regulatory Networks. PLOS ONE 10, e0133387. 10.1371/journal.pone.0133387

Diaz de la Loza and Thompson 2017. Forces shaping the Drosophila wing. Mechanisms of Development, 23 Oct 2016, 144(Pt A):23–32. DOI: 10.1016/j.mod.2016.10.003

Drummond-Barbosa and Spradling 2001. Stem cells and their progeny respond to nutritional changes during Drosophila oogenesis. Dev Biol 231: 265–278. 10.1006/dbio.2000.0135

Evans, C.J., Olson, J.M., Ngo, K.T., Kim, E., Lee, N.E., Kuoy, E., Patananan, A.N., Sitz, D., Tran, P., Do, M.-T., Yackle, K., Cespedes, A., Hartenstein, V., Call, G.B., Banerjee, U., 2009. G-TRACE: rapid Gal4-based cell lineage analysis in Drosophila. Nat. Methods 6, 603–605. 10.1038/nmeth.1356

Fox, D.T., Gall, J.G., Spradling, A.C., 2010. Error-prone polyploid mitosis during normal Drosophila development. Genes Dev. 24, 2294–2302. 10.1101/gad.1952710

Friard, O., Gamba, M., 2016. BORIS: a free, versatile open-source event-logging software for video/audio coding and live observations. Methods Ecol. Evol. 7, 1325–1330. 10.1111/2041-210X.12584

Gohl, D., Müller, M., Pirrotta, V., Affolter, M., Schedl, P., 2008. Enhancer blocking and transvection at the Drosophila apterous locus. Genetics 178, 127–143. 10.1534/genetics.107.077768

Golic, K.G., Golic, M.M., 1996. Engineering the Drosophila genome: chromosome rearrangements by design. Genetics 144, 1693–1711. 10.1093/genetics/144.4.1693

Gratz, S.J., Cummings, A.M., Nguyen, J.N., Hamm, D.C., Donohue, L.K., Harrison, M.M., Wildonger, J., O’connor-Giles, K.M., 2013. Genome engineering of Drosophila with the CRISPR RNA-guided Cas9 nuclease. Genetics 194, 1029–1035. 10.1534/genetics.113.152710

Hadjieconomou, D., King, G., Gaspar, P., Mineo, A., Blackie, L., Ameku, T., Studd, C., de Mendoza, A., Diao, F., White, B.H., Brown, A.E.X., Plaçais, P.-Y., Préat, T., Miguel-Aliaga, I., 2020. Enteric neurons increase maternal food intake during reproduction. Nature 587, 455–459. 10.1038/s41586-020-2866-8

Herzig, M.C., Thor, S., Thomas, J.B., Reichert, H., Hirth, F., 2001. Expression and function of the LIM homeodomain protein Apterous during embryonic brain development of Drosophila. Dev. Genes Evol. 211, 545–554. 10.1007/s00427-001-0195-y

Hinnant et al 2020. Coordinating Proliferation, Polarity, and Cell Fate in the *Drosophila* Female Germline. Front Cell Dev Biol Feb 4:8:19. DOI: 10.3389/fcell.2020.00019

Irvine, K.D., Rauskolb, C., 2001. Boundaries in development: formation and function. Annu. Rev. Cell Dev. Biol. 17, 189–214. 10.1146/annurev.cellbio.17.1.189

Itskov, P.M., Moreira, J.-M., Vinnik, E., Lopes, G., Safarik, S., Dickinson, M.H., Ribeiro, C., 2014. Automated monitoring and quantitative analysis of feeding behaviour in Drosophila. Nat. Commun. 5, 4560. 10.1038/ncomms5560

Iwaki and Lengyel 2002. A Delta-Notch signaling border regulated by Engrailed/Invected repression specifies boundary cells in the Drosophila hindgut. Mech. Dev. 114:71—84. 10.1016/s0925-4773(02)00061-8

Keles, et al., 2025. FlyVISTA, an integrated machine learning platform for deep phenotyping of sleep in Drosophila. Sci. Adv. 11(11), eadq8131. 10.1126/sciadv.adq8131

King and Sang 1958. Additional description of ap^4^. Drosophila Inform. Serv. 32:133.

King and Bodenstein 1965. The transplantation of ovaries between genetically sterile and wild-type Drosophila melanogaster. Z. Naturforsch. 20b: 292–297. 10.1515/znb-1965-0404

Lemaitre and Miguel-Aliaga 2023. The Digestive Tract of Drosophila melanogaster. Annual Review of Genetics 47:377–404. DOI: 10.1146/annurev-genet-111212-133343

Linford, N. J., Bilgir, C., Ro, J., & Pletcher, S. D. (2013). Measurement of lifespan in Drosophila melanogaster. Journal of visualized experiments: JoVE, (71), 50068. 10.3791/50068

Liu, Q., Yang, X., Luo, M., Su, J., Zhong, J., Li, X., Chan, R.H.M., Wang, L., 2023. An iterative neural processing sequence orchestrates feeding. Neuron 111, 1651–1665.e5. 10.1016/j.neuron.2023.02.025

Lundgren et al 1995. Control of neuronal pathway selection by the *Drosophila* LIM homeodomain gene *apterous*. Development 121, 1769–1773. 10.1242/dev.121.6.1769

Marianes and Spradling 2013. Physiological and stem cell compartmentalization within the Drosophila midgut. eLife 2013;2:e00886

Magwere et al 2004. Sex Differences in the Effect of Dietary Restriction on Life Span and Mortality Rates in Female and Male *Drosophila Melanogaster*. The Journals of Gerontology: Series A, Volume 59, Issue 1, Pages B3–B9. 10.1093/gerona/59.1.B3

McGuire, S.E., Mao, Z., Davis, R.L., 2004. Spatiotemporal gene expression targeting with the TARGET and gene-switch systems in Drosophila. Sci. STKE Signal Transduct. Knowl. Environ. 2004, pl6. 10.1126/stke.2202004pl6

Medina et al 2022. Investigating local and systemic intestinal signalling in health and disease with *Drosophila*. Dis Model Mech (2022) 15 (3): dmm049332. 10.1242/dmm.049332

Meiselman, M.R., Kingan, T.G., Adams, M.E., 2018. Stress-induced reproductive arrest in Drosophila occurs through ETH deficiency-mediated suppression of oogenesis and ovulation. BMC Biol. 16, 18. 10.1186/s12915-018-0484-9

Messina, M., Angotti, R., Molinaro, F., 2016. Meconium Plug Syndrome, in: Buonocore, G., Bracci, R., Weindling, M. (Eds.), Neonatology: A Practical Approach to Neonatal Diseases. Springer International Publishing, Cham, pp. 1–6. 10.1007/978-3-319-18159-2_231-1

Miguel-Aliaga, et al., 2018. Anatomy and Physiology of the Digestive Tract of *Drosophila melanogaster*. Genetics 210(2), 357–396, 10.1534/genetics.118.300224

Milán, M. et al., 1998. *Beadex* encodes an LMO protein that regulates Apterous LIM– homeodomain activity in *Drosophila* wing development: a model for *LMO* oncogene function. Genes Dev 12(18):2912–2920. doi: 10.1101/gad.12.18.2912

Milán, M., Cohen, S.M., 2000. Temporal regulation of apterous activity during development of the Drosophila wing. Development 127(14):3069–78. doi: 10.1242/dev.127.14.3069.

Metz, C.W., 1914. An Apterous Drosophila and Its Genetic behaviour. Am. Nat. 48, 675–692. 10.1086/279439

Müller, B., Basler, K., 2000. The repressor and activator forms of Cubitus interruptus control Hedgehog target genes through common generic gli-binding sites. Dev. Camb. Engl. 127, 2999–3007. 10.1242/dev.127.14.2999

Murakami, R., Shigenaga, A., Kawakita, M., Takimoto, K., Yamaoka, I., Akasaka, K., Shimada, H., 1995. aproctous, a locus that is necessary for the development of the proctodeum in Drosophila embryos, encodes a homolog of the vertebrate Brachyury gene. Rouxs Arch. Dev. Biol. 205, 89–96. 10.1007/BF00188847

O’Brien et al 2011. Altered Modes of Stem Cell Division Drive Adaptive Intestinal Growth. Cell Volume 147, Issue 3, Pages 603–614. 10.1016/j.cell.2011.08.048

O’Keefe, D.D., Thor, S., Thomas, J.B., 1998. Function and specificity of LIM domains in Drosophila nervous system and wing development. Dev. Camb. Engl. 125, 3915–3923. 10.1242/dev.125.19.3915

Pereira, T.D., Tabris, N., Matsliah, A., Turner, D.M., Li, J., Ravindranath, S., Papadoyannis, E.S., Normand, E., Deutsch, D.S., Wang, Z.Y., McKenzie-Smith, G.C., Mitelut, C.C., Castro, M.D., D’Uva, J., Kislin, M., Sanes, D.H., Kocher, S.D., Wang, S.S.-H., Falkner, A.L., Shaevitz, J.W., Murthy, M., 2022. SLEAP: A deep learning system for multi-animal pose tracking. Nat. Methods 19, 486–495. 10.1038/s41592-022-01426-1

Plusczyk, T., Bolli, M., Schilling, M., 2006. [Ileus disease]. Chir. Z. Alle Geb. Oper. Medizen 77, 898–903. 10.1007/s00104-006-1237-9

Pfeiffer, B.D., Jenett, A., Hammonds, A.S., Ngo, T.-T.B., Misra, S., Murphy, C., Scully, A., Carlson, J.W., Wan, K.H., Laverty, T.R., Mungall, C., Svirskas, R., Kadonaga, J.T., Doe, C.Q., Eisen, M.B., Celniker, S.E., Rubin, G.M., 2008. Tools for neuroanatomy and neurogenetics in Drosophila. Proc. Natl. Acad. Sci. U. S. A. 105, 9715–9720. 10.1073/pnas.0803697105

Postlethwait, J.H., Weiser, K., 1973. Vitellogenesis induced by Juvenile Hormone in the Female Sterile Mutant apterous-four in Drosophila melanogaster. Nature. New Biol. 244, 284–285. 10.1038/newbio244284a0

Rakestraw, P.C., Hardy, J., 2006. Chapter 36 - Large Intestine, in: Auer, J.A., Stick, J.A. (Eds.), Equine Surgery (Third Edition). W.B. Saunders, Saint Louis, pp. 436–478. 10.1016/B1-41-600123-9/50038-3

Regan et al 2016. Sex difference in pathology of the ageing gut mediates the greater response of female lifespan to dietary restriction. eLife 10.7554/eLife.10956

Richard, D.S., Arnim, A.E., Gilbert, L.I., 1993. A reappraisal of the hormonal regulation of larval fat body histolysis in female Drosophila melanogaster. Experientia 49, 150–156. 10.1007/BF01989420

Sathe, M., Houwen, R., 2017. Meconium ileus in Cystic Fibrosis. J. Cyst. Fibros. Off. J. Eur. Cyst. Fibros. Soc. 16 Suppl 2, S32–S39. 10.1016/j.jcf.2017.06.007

Schoborg, T.A., Smith, S.L., Smith, L.N., Morris, H.D., Rusan, N.M., 2019. Micro-computed tomography as a platform for exploring Drosophila development. Dev. Camb. Engl. 146. 10.1242/dev.176685

Schoenfelder, K.P., Montague, R.A., Paramore, S.V., Lennox, A.L., Mahowald, A.P., Fox, D.T., 2014. Indispensable pre-mitotic endocycles promote aneuploidy in the Drosophila rectum. Development 141, 3551–3560. 10.1242/dev.109850

Schwenke, R.A., Lazzaro, B.P., Wolfner, M.F., 2016. Reproduction-Immunity Trade-Offs in Insects. Annu. Rev. Entomol. 61, 239–256. 10.1146/annurev-ento-010715-023924

Shaw, P.J., Cirelli, C., Greenspan, R.J., Tononi, G., 2000. Correlates of Sleep and Waking in Drosophila melanogaster. Science 287, 1834–1837. 10.1126/science.287.5459.1834

Stevens, M.E., Bryant, P.J., 1986. TEMPERATURE-DEPENDENT EXPRESSION OF THE APTEROUS PHENOTYPE IN DROSOPHILA MELANOGASTER. Genetics 112, 217–228. 10.1093/genetics/112.2.217

Stevens, M.E., Bryant, P.J., 1985. Apparent genetic complexity generated by developmental thresholds: the apterous locus in Drosophila melanogaster. Genetics 110, 281–297. 10.1093/genetics/110.2.281

Stratmann, J., Thor, S., 2017. Neuronal cell fate specification by the molecular convergence of different spatio-temporal cues on a common initiator terminal selector gene. PLOS Genet. 13, e1006729. 10.1371/journal.pgen.1006729

Takashima, S., Mkrtchyan, M., Younossi-Hartenstein, A., Merriam, J.R., Hartenstein, V., 2008. The behaviour of Drosophila adult hindgut stem cells is controlled by Wnt and Hh signalling. Nature 454, 651–655. 10.1038/nature07156

Thibault, S.T., Singer, M.A., Miyazaki, W.Y., Milash, B., Dompe, N.A., Singh, C.M., Buchholz, R., Demsky, M., Fawcett, R., Francis-Lang, H.L., Ryner, L., Cheung, L.M., Chong, A., Erickson, C., Fisher, W.W., Greer, K., Hartouni, S.R., Howie, E., Jakkula, L., Joo, D., Killpack, K., Laufer, A., Mazzotta, J., Smith, R.D., Stevens, L.M., Stuber, C., Tan, L.R., Ventura, R., Woo, A., Zakrajsek, I., Zhao, L., Chen, F., Swimmer, C., Kopczynski, C., Duyk, G., Winberg, M.L., Margolis, J., 2004. A complementary transposon tool kit for Drosophila melanogaster using P and piggyBac. Nat. Genet. 36, 283–287. 10.1038/ng1314

Tomita, T., Nakamura, M., Kobayashi, Y., Yoshinaka, A., Murakumo, K., 2020. Viviparous stingrays avoid contamination of the embryonic environment through faecal accumulation mechanisms. Sci. Rep. 10, 7378. 10.1038/s41598-020-64271-2

van Alphen, B., Semenza, E.R., Yap, M., van Swinderen, B., Allada, R., 2021. A deep sleep stage in Drosophila with a functional role in waste clearance. Sci. Adv. 7. 10.1126/sciadv.abc2999

Waldhausen, J.H.T., Richards, M., 2018. Meconium Ileus. Clin. Colon Rectal Surg. 31, 121–126. 10.1055/s-0037-1609027

Wang, Y., Li, W., Zhou, C., Zhao, Zifeng, Ma, J., Jiang, H., Wei, M., Gao, Y., Dai, Y., Zhang, X., Yang, N., Feng, F., Zhang, J., Ji, Y., Liu, J., Zhang, C., Li, L., Jiang, X., Li, Z., Zhao, Zengren, 2024. Mortality risk of patients with intestinal obstruction. BMC Cancer 24, 1062. 10.1186/s12885-024-12834-1

Wessing, A., Eichelberg, D., 1973. Elektronenmikroskopische Untersuchungen zur Struktur und Funktion der Rektalpapillen von Drosophila melanogaster. Z. Für Zellforsch. Mikrosk. Anat. 136, 415–432. 10.1007/BF00307043

Wilson, T. G., 1981a. Expression of phenotypes in a temperature-sensitive allele of the apterous mutation in Drosophila melanogaster. Dev. Biol. 85, 425–433. 10.1016/0012-1606(81)90274-8

Wilson, T. G., 1981b. A mosaic analysis of the apterous mutation in Drosophila melanogaster. Dev. Biol. 85, 434–445. 10.1016/0012-1606(81)90275-x

Xiao, W., Jiao, Z.-L., Senol, E., Yao, J., Zhao, M., Zhao, Z.-D., Chen, X., Cao, P., Fu, Y., Gao, Z., Shen, W.L., Xu, X.-H., 2022. Neural circuit control of innate behaviors. Sci. China Life Sci. 65, 466–499. 10.1007/s11427-021-2043-2

Youn, H., Kirkhart, C., Chia, J., Scott, K., 2018. A subset of octopaminergic neurons that promotes feeding initiation in Drosophila melanogaster. PLOS ONE 13, e0198362. 10.1371/journal.pone.0198362

