## Supplementary figures and images for "Intestinal control of feeding initiation in *Drosophila melanogaster*"

### Supplementary Figure 1

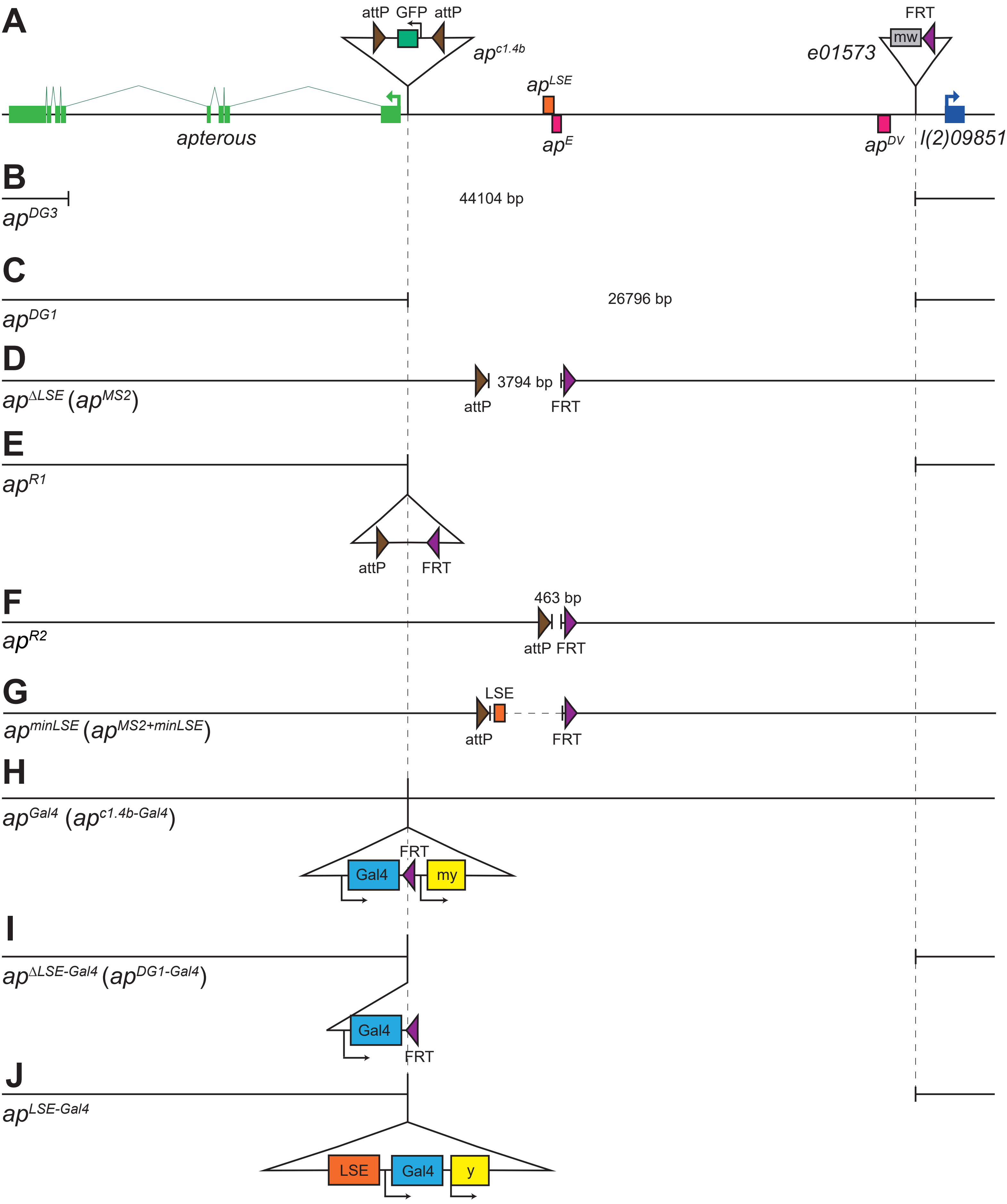


**Reinger et al. Suppl Figure 1**

### Supplementary Figure 2

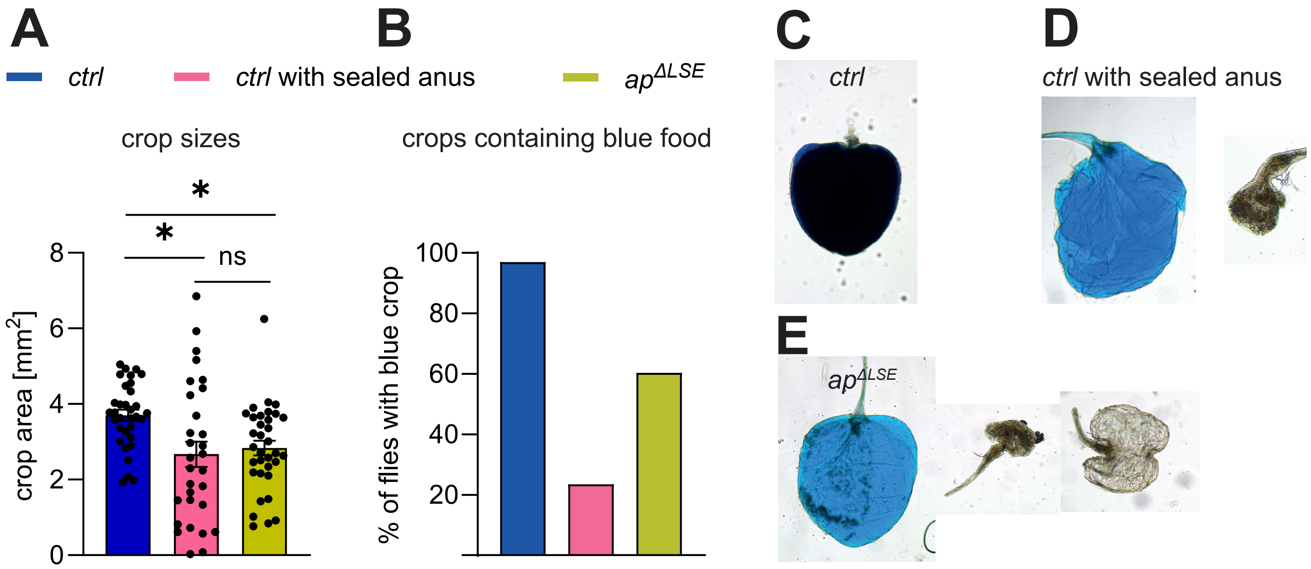


**Reinger et al. Suppl Figure 2**

### Supplementary Figure 3

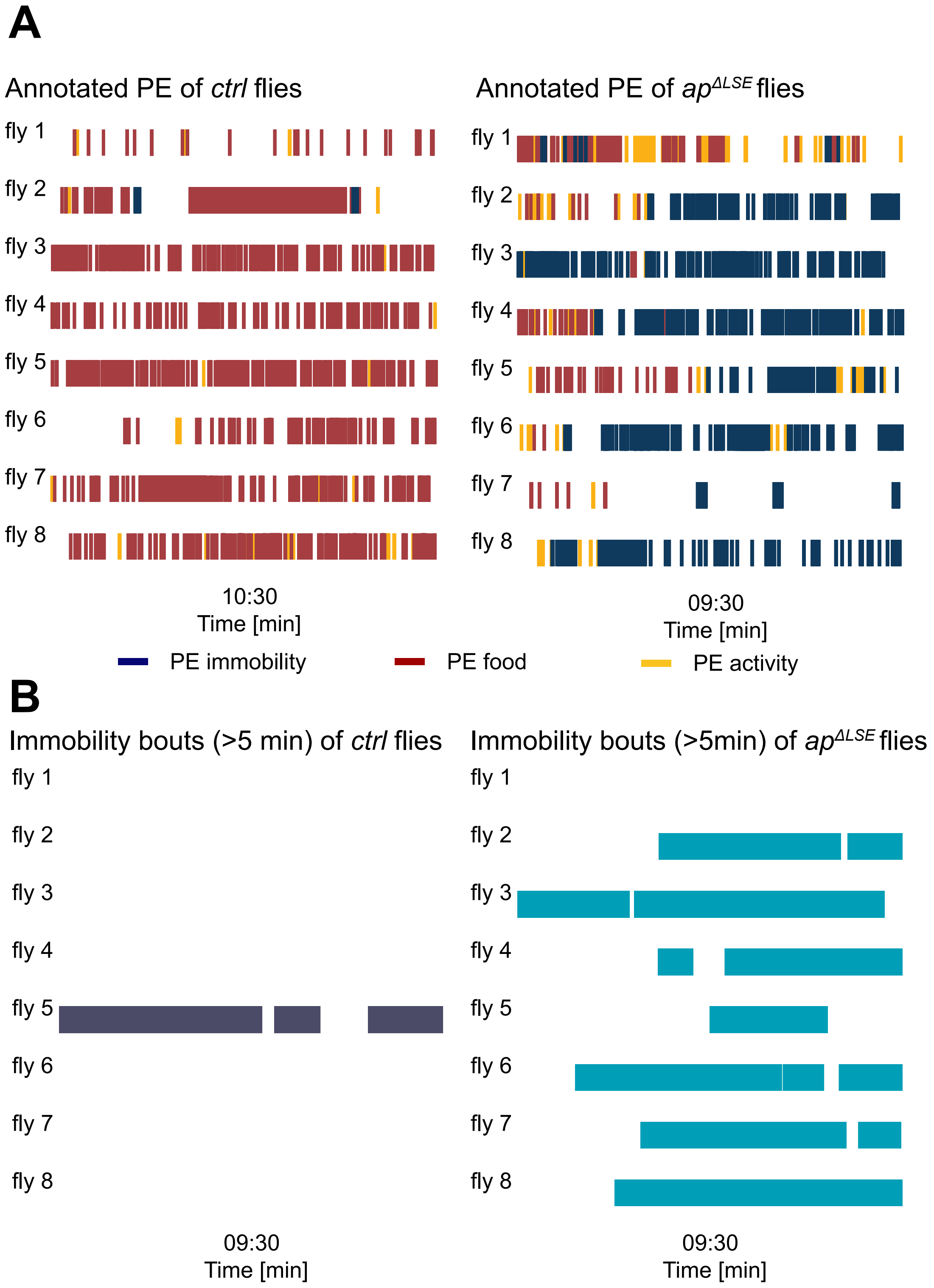


**Reinger et al. Suppl Figure 3**

### Supplementary Figure 4

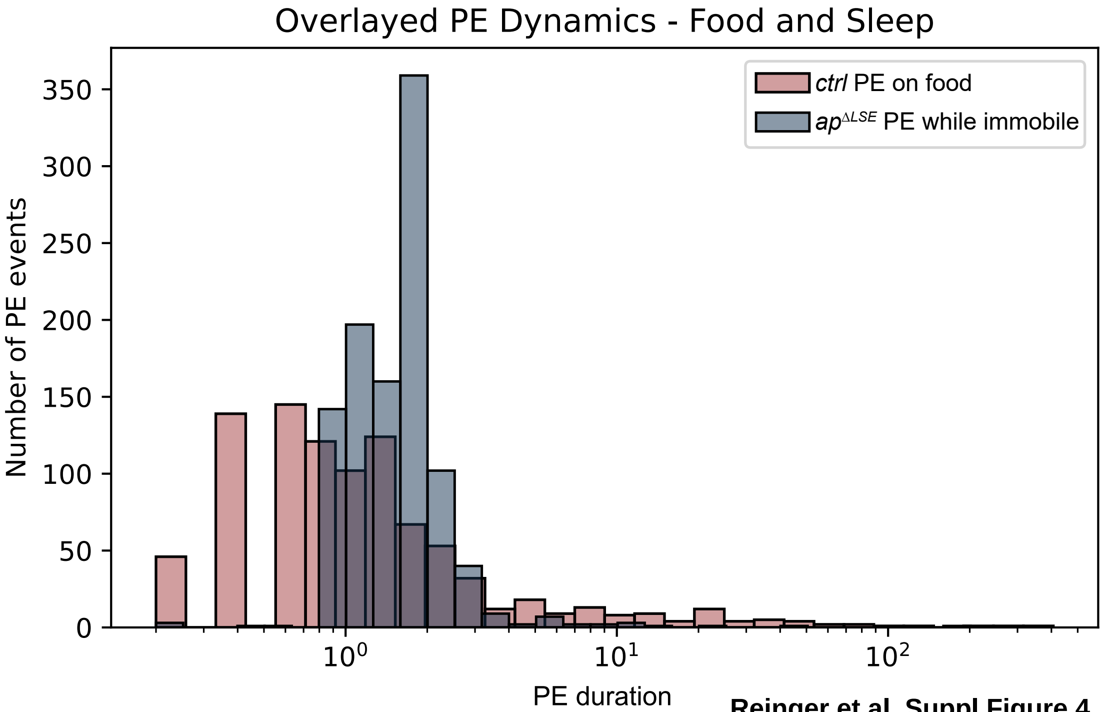


**Reinger et al. Suppl Figure 4**

### Supplementary Figure 5

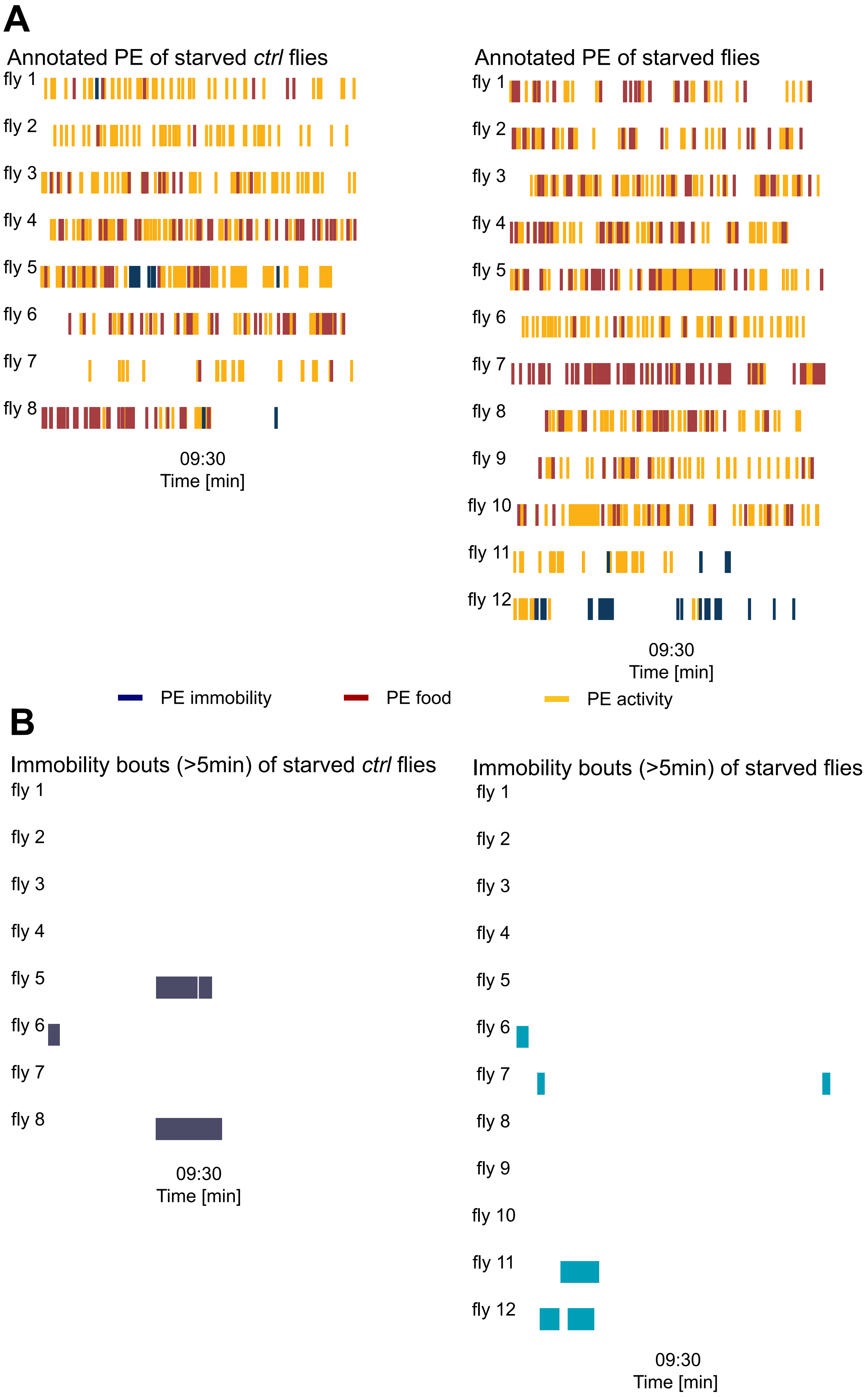


**Reinger et al. Suppl Figure 5**

### Supplementary Figure 6

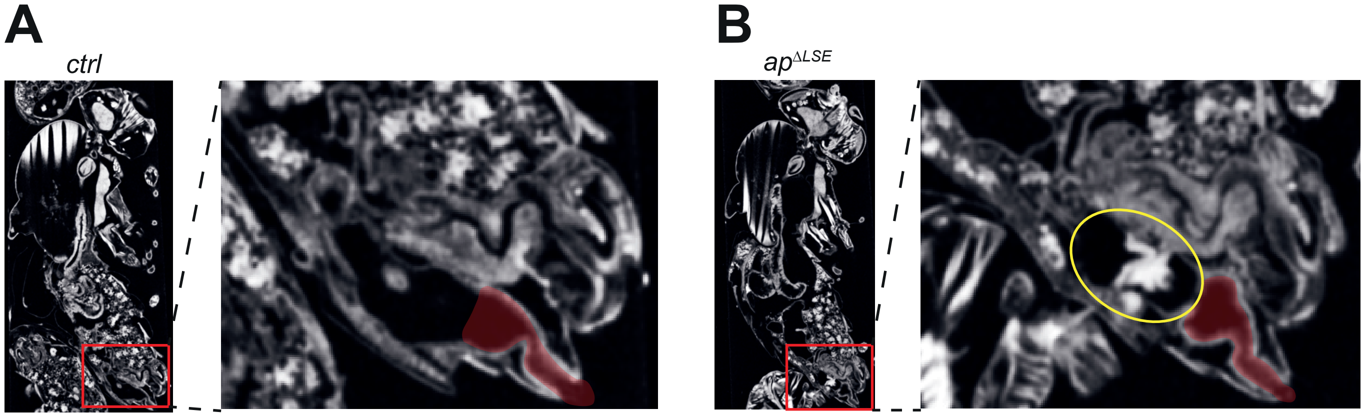


**Reinger et al. Suppl Figure 6**

### Supplementary Figure 7

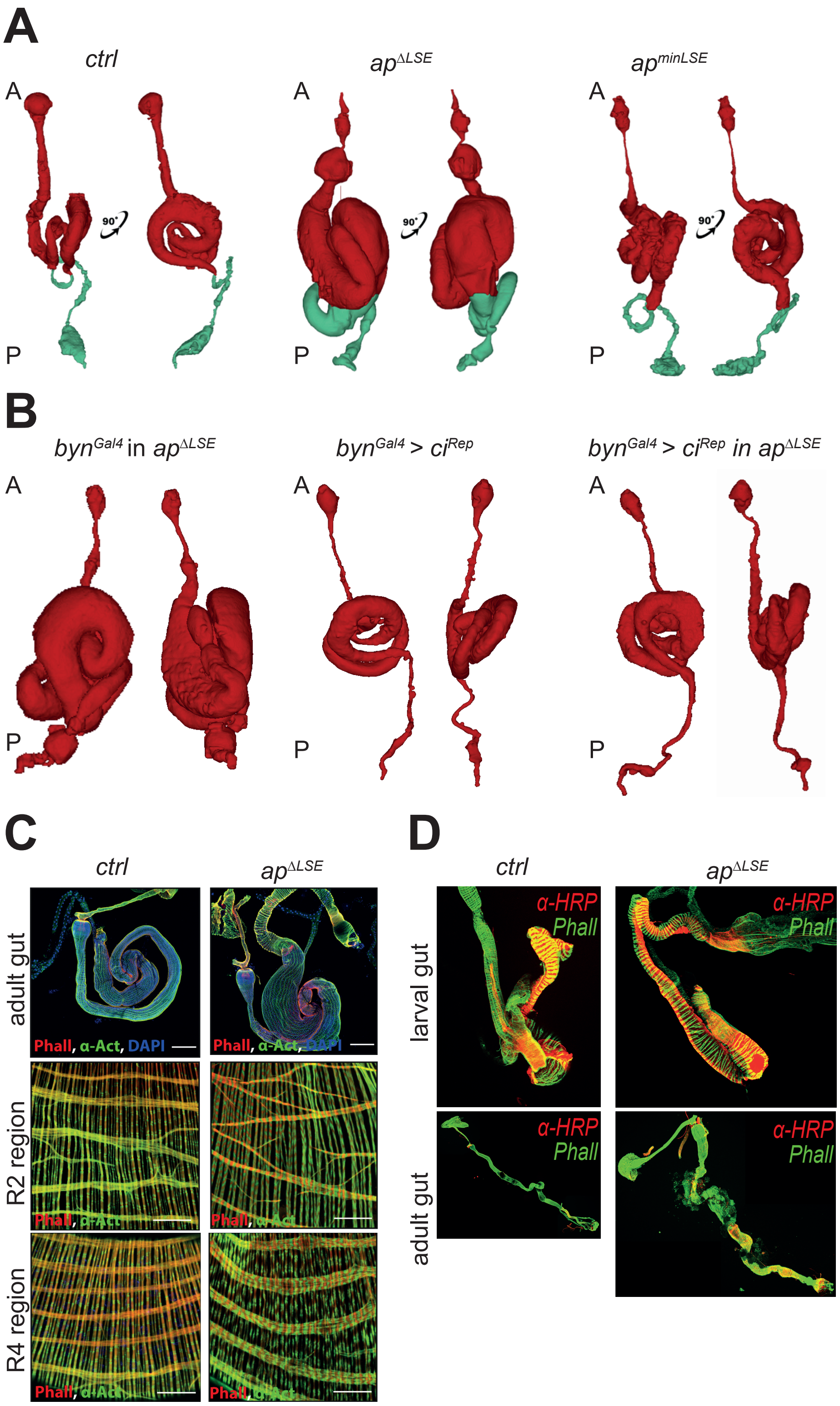


**Reinger et al. Suppl Figure 7**

### Supplementary Figure 8

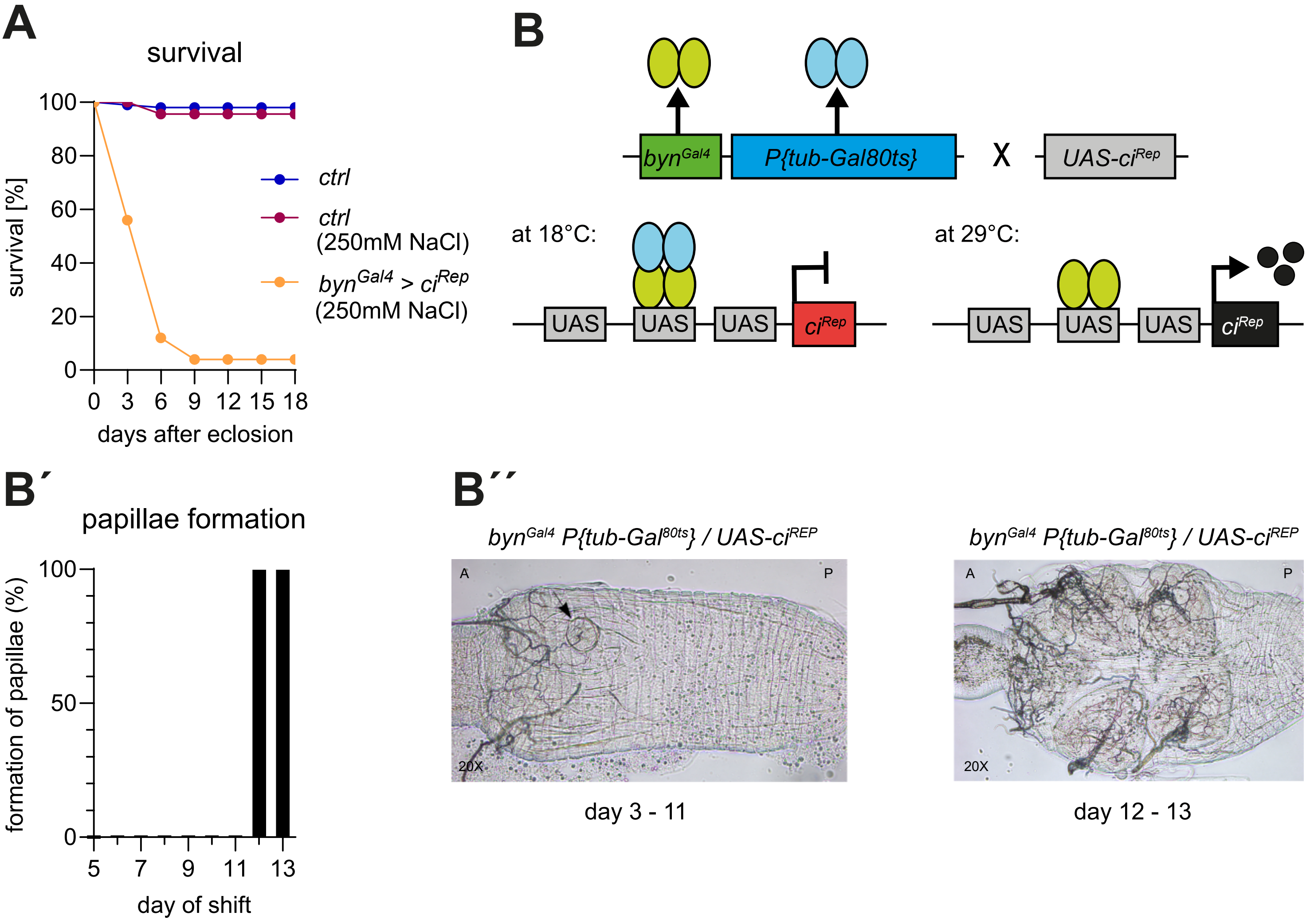


**Reinger et al. Suppl Figure 8**

### Supplementary Figure 9

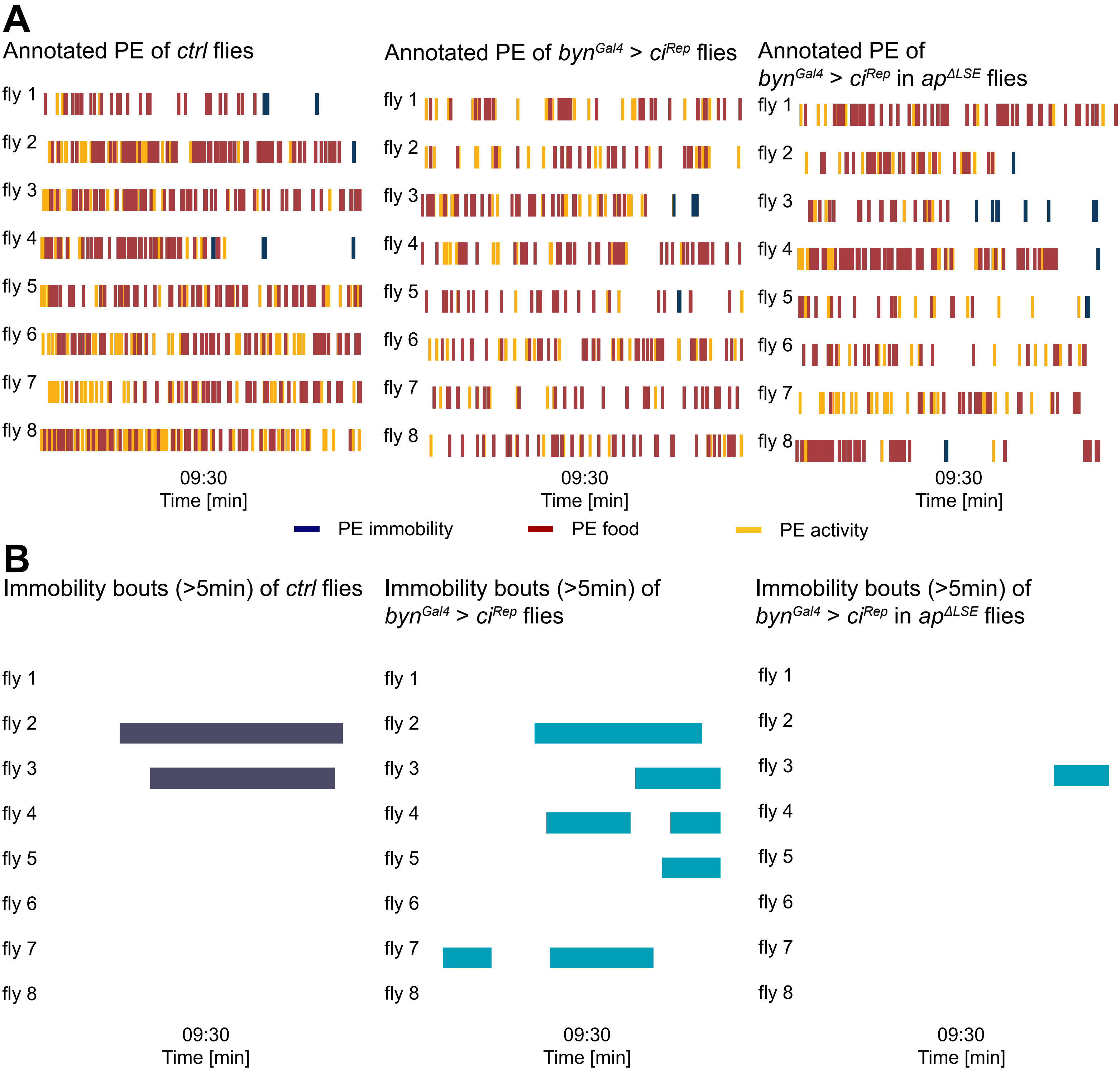


**Reinger et al. Suppl Figure 9**

### Supplementary Figure 10

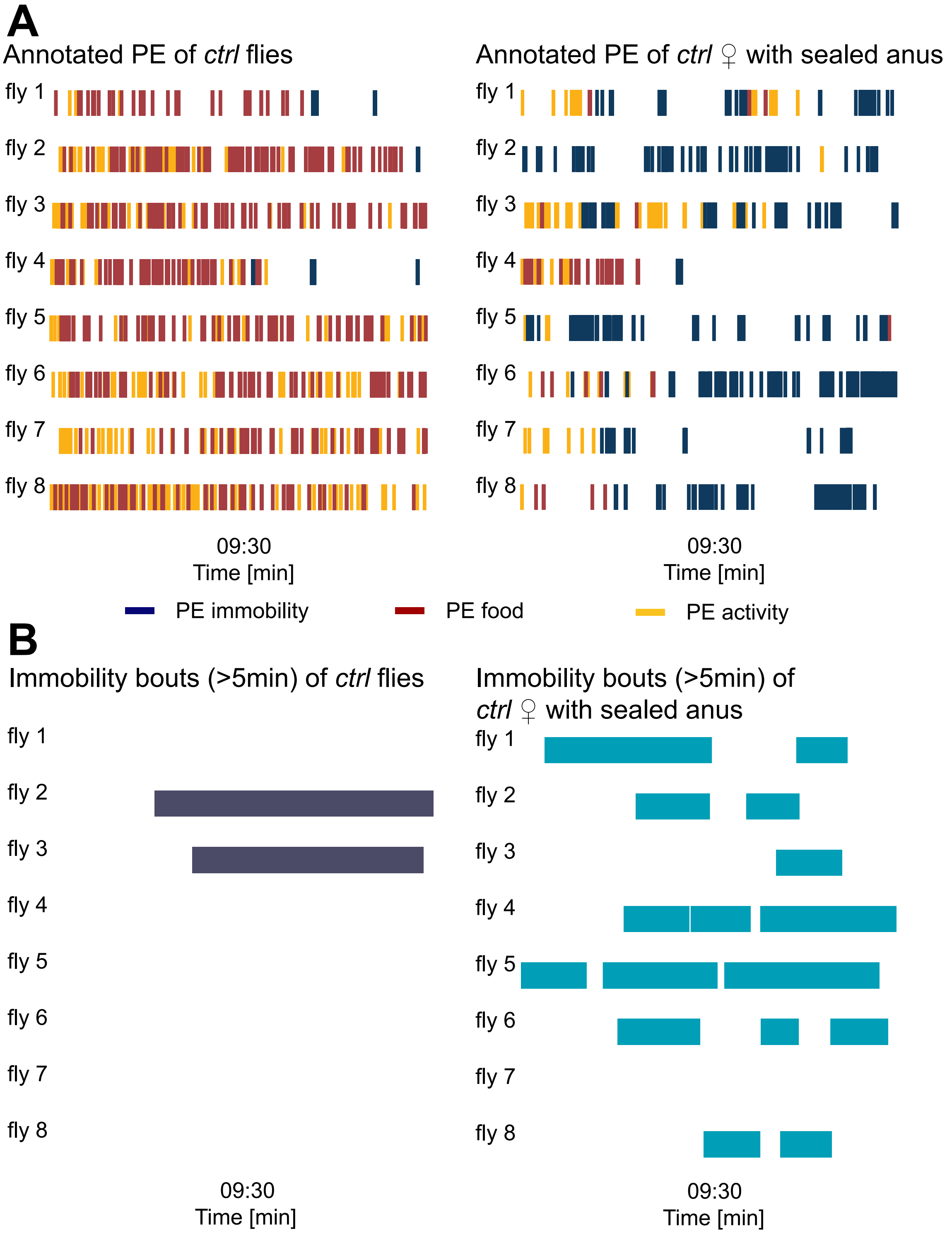


**Reinger et al. Suppl Figure 10**
