## Supplementary Table 1 for "Intestinal control of feeding initiation in *Drosophila melanogaster*"

### Supplementary Table 1. Statistical analyses.

| Figure | Statistical test | Pairwise comparison | Test statistic | *P* |
|---|---|---|---|---|
| **1b** | Mann-Whitney test | | | |
| | *ctrl* females | *ctrl* females vs *ctrl* males | U=22 | 0.7851 |
| | *ctrl* males | | | |
| **1c** | Mann-Whitney test | *ctrl* females vs. *ctrl* males | U=45.5 | 0.1239 |
| | *ctrl* females | | | |
| | *ctrl* males | | | |
| **1e** | Mann-Whitney test | *ctrl* females (+food) vs. *ctrl* females (-food) | U=3.5 | 0.2286 |
| | *ctrl* females (+food) | *ctrl* males (+food) vs. *ctrl* males (-food) | U=3.5 | 0.2286 |
| | *ctrl* females (-food) | | | |
| | *ctrl* males (+food) | | | |
| | *ctrl* males (-food) | | | |
| **2c** | Kruskal Wallis ANOVA | | | |
| | Dunns post-hoc test | *ctrl vs. apΔLSE / +* | z=1.485 | 0.8253 |
| | Dunns post-hoc test | *ctrl vs. apΔLSE* | z=1.428 | 0.9200 |
| | Dunns post-hoc test | *ctrl vs. apminLSE* | z=0.3398 | >0.9999 |
| **2d** | Kruskal Wallis ANOVA | | | |
| | Dunns post-hoc test | *ctrl* vs. $ap^{\Delta LSE}$ / + | z=1.568 | 0.7016 |
| | Dunns post-hoc test | *ctrl* vs. $ap^{\Delta LSE}$ | z=2.679 | 0.0420 |
| | Dunns post-hoc test | *ctrl* vs. $ap^{minLSE}$ | z=0.07118 | >0.9999 |
| **2e** | Kruskal Wallis ANOVA | | | |
| | Dunns post-hoc test | *ctrl* vs. $ap^{\Delta LSE}$ / + | z=0.5921 | >0.9999 |
| | Dunns post-hoc test | *ctrl* vs. $ap^{\Delta LSE}$ | z=6.577 | <0.0001 |
| | Dunns post-hoc test | *ctrl* vs. $ap^{minLSE}$ | z=3.265 | 0.0066 |
| **2g** | Mann-Whitney test | | | |
| | *ctrl* females | *ctrl* females vs. $ap^{\Delta LSE}$ females on food | U=0 | 0.0002 |
| | $ap^{\Delta LSE}$ females | *ctrl* females vs. $ap^{\Delta LSE}$ females while active | U=13 | 0.0499 |
| | | *ctrl* females vs. $ap^{\Delta LSE}$ females while immobile | U=0 | 0.0002 |
| **2h** | Mann-Whitney test | | | |
| | *ctrl* females | *ctrl* females vs. $ap^{\Delta LSE}$ females activity | U=8.5 | 0.0064 |
| | $ap^{\Delta LSE}$ females | *ctrl* females vs. $ap^{\Delta LSE}$ females immobility | U=8.5 | 0.0064 |
| **2i** | Mann-Whitney test | | | |
| | *ctrl* females | *ctrl* females vs. $ap^{\Delta LSE}$ females | U=8.5 | 0.0064 |
| | $ap^{\Delta LSE}$ females | | | |
| **2j** | Mann-Whitney test | | | |
| | *ctrl* females (fed) | *ctrl* females (fed) vs. *ctrl* females (starved) on food | U=38 | 0.4727 |
| | *ctrl* females (starved) | *ctrl* females (fed) vs. *ctrl* females (starved) while active | U=44 | 0.7921 |
| | | *ctrl* females (fed) vs. *ctrl* females (starved) while immobile | U=40 | 0.5788 |
| **2k** | Mann-Whitney test | | | |
| | *ctrl* females (fed) | *ctrl* females (fed) vs. *ctrl* females (starved) activity | U=46.50 | >0.9999 |
| | *ctrl* females (starved) | *ctrl* females (fed) vs. *ctrl* females (starved) immobility | U=46.50 | >0.9999 |
| **5d** | Kruskal Wallis ANOVA | | | |
| | Dunns post-hoc test | *ctrl* vs. $byn^{Gal4}$ | z=1.505 | 0.7938 |
| | Dunns post-hoc test | *ctrl* vs. $byn^{Gal4}$ *in* $ap^{\Delta LSE}$ | z=2.614 | 0.0537 |
| | Dunns post-hoc test | *ctrl* vs. $byn^{Gal4}$ > $ci^{Rep}$ *in* $ap^{\Delta LSE}$ | z=0.1058 | >0.9999 |
| **5e** | Kruskal Wallis ANOVA | | | |
| | Dunns post-hoc test | *ctrl* vs. $byn^{Gal4}$ > $ci^{Rep}$ on food | z=1.595 | 0.3348 |
| | Dunns post-hoc test | *ctrl* vs. $byn^{Gal4}$ > $ci^{Rep}$ *in* $ap^{\Delta LSE}$ on food | z=1.595 | 0.3348 |
| | Dunns post-hoc test | $byn^{Gal4}$ > $ci^{Rep}$ vs. $byn^{Gal4}$ > $ci^{Rep}$ *in* $ap^{\Delta LSE}$ on food | z=0 | >0.9999 |
| | Dunns post-hoc test | *ctrl* vs. $byn^{Gal4}$ > $ci^{Rep}$ while active | z=1.857 | 0.1901 |
| | Dunns post-hoc test | *ctrl* vs. $byn^{Gal4}$ > $ci^{Rep}$ *in* $ap^{\Delta LSE}$ while active | z=2.016 | 0.1315 |
| | Dunns post-hoc test | $byn^{Gal4}$ > $ci^{Rep}$ vs. $byn^{Gal4}$ > $ci^{Rep}$ *in* $ap^{\Delta LSE}$ while active | z=0.1591 | >0.9999 |
| | Dunns post-hoc test | *ctrl* vs. $byn^{Gal4}$ > $ci^{Rep}$ while immobile | z=0.4550 | >0.9999 |
| | Dunns post-hoc test | *ctrl* vs. $byn^{Gal4}$ > $ci^{Rep}$ *in* $ap^{\Delta LSE}$ while immobile | z=0.7500 | >0.9999 |
| | Dunns post-hoc test | $byn^{Gal4}$ > $ci^{Rep}$ vs. $byn^{Gal4}$ > $ci^{Rep}$ *in* $ap^{\Delta LSE}$ while immobile | z=1.204 | 0.6858 |
| **5f** | Kruskal Wallis ANOVA | | | |
| | Dunns post-hoc test | *ctrl* vs. $byn^{Gal4}$ > $ci^{Rep}$ activity | z=0.9057 | >0.9999 |
| | Dunns post-hoc test | *ctrl* vs. $byn^{Gal4}$ > $ci^{Rep}$ *in* $ap^{\Delta LSE}$ activity | z=1.158 | 0.7400 |
| | Dunns post-hoc test | $byn^{Gal4}$ > $ci^{Rep}$ vs. $byn^{Gal4}$ > $ci^{Rep}$ *in* $ap^{\Delta LSE}$ activity | z=2.064 | 0.1170 |
| | Dunns post-hoc test | *ctrl* vs. $byn^{Gal4}$ > $ci^{Rep}$ while immobility | z=0.9057 | >0.9999 |
| | Dunns post-hoc test | *ctrl* vs. $byn^{Gal4}$ > $ci^{Rep}$ *in* $ap^{\Delta LSE}$ immobility | z=1.158 | 0.7400 |
| | Dunns post-hoc test | $byn^{Gal4}$ > $ci^{Rep}$ vs. $byn^{Gal4}$ > $ci^{Rep}$ *in* $ap^{\Delta LSE}$ immobility | z=2.064 | 0.1170 |
| **5g** | Kruskal Wallis ANOVA | | | |
| | Dunns post-hoc test | *ctrl* vs. $byn^{Gal4}$ > $ci^{Rep}$ | z=0.9057 | >0.9999 |
| | Dunns post-hoc test | *ctrl* vs. $byn^{Gal4}$ > $ci^{Rep}$ *in* $ap^{\Delta LSE}$ | z=1.158 | 0.7400 |
| | Dunns post-hoc test | $byn^{Gal4}$ > $ci^{Rep}$ vs. $byn^{Gal4}$ > $ci^{Rep}$ *in* $ap^{\Delta LSE}$ | z=2.064 | 0.1170 |
| **Sup 2a** | Kruskal Wallis ANOVA | | | |
| | Dunns post-hoc test | *ctrl* vs. $ap^{\Delta LSE}$ | z=3.245 | 0.0035 |
| | Dunns post-hoc test | *ctrl* vs. *ctrl* with sealed anus | z=2.952 | 0.0095 |
| | Dunns post-hoc test | *ctrl* with sealed anus vs. $ap^{\Delta LSE}$ | z=0.3888 | >0.9999 |
