## Supplementary Table 2 for "Intestinal control of feeding initiation in *Drosophila melanogaster*"

### Supplementary Table 2. Detailed description of genotypes.

| Figure | Name in Figure | genotype |
|---|---|---|
| **1b-f** | *ctrl* | *y w* |
| **2b** | *ctrl*<br>$ap^{\Delta LSE}$ / +<br>$ap^{\Delta LSE}$<br>$ap^{minLSE}$ | *y w*<br>*y w* ; $ap^{DG3}$ / +<br>*y w* ; $ap^{DG3}$ / $ap^{MS2}$<br>*y w* ; $ap^{MS2+minLSE}$ / $ap^{DG3}$ |
| **2c-e** | *ctrl*<br>$ap^{\Delta LSE}$ / +<br>$ap^{\Delta LSE}$<br>$ap^{minLSE}$ | *y w*<br>*y w* ; $ap^{DG3}$ / +<br>*y w* ; $ap^{DG3}$ / $ap^{MS2}$<br>*y w* ; $ap^{MS2+minLSE}$ / $ap^{MS2}$ |
| **2f-k** | *ctrl*<br>$ap^{\Delta LSE}$ | *y w*<br>*y w* ; $ap^{DG3}$ / $ap^{MS2}$ |
| **3a** | $byn^{Gal4}$ > $mCherry^{NLS}$ | *y w* ; + / + ; $byn^{Gal4}$ / *UAS*-$mCherry^{NLS}$ |
| **3b** | $ap^{Gal4}$ > *G-TRACE*<br>$ap^{\Delta LSE\text{-}Gal4}$ > *G-TRACE*<br>$ap^{minLSE\text{-}Gal4}$ > *G-TRACE* | *y w* ; $ap^{c1.4b\text{-}Gal4}$ / $ap^{DG3}$ ; *G-TRACE* / +<br>*y w* ; $ap^{DG1\text{-}Gal4}$ / $ap^{DG3}$ ; *G-TRACE* / +<br>*y w* ; $ap^{LSE\text{-}Gal4}$ / $ap^{DG3}$ ; *G-TRACE* / + |
| **3c** | *ctrl*<br>$ap^{\Delta LSE}$<br>$ap^{minLSE}$ | *y w*<br>*y w* ; $ap^{DG3}$ / $ap^{MS2}$<br>*y w* ; $ap^{MS2+minLSE}$ / $ap^{MS2}$ |
| **4a-a´** | *ctrl*<br>$Dl^{Gal4}$ >$mCherry^{NLS}$ in *ctrl* | *y w*<br>*y w* ; $ap^{DG3}$ / + ; $Dl^{Gal4}$ / *UAS*-$mCherry^{NLS}$ |
| **4b-b´** | $ap^{\Delta LSE}$<br>$Dl^{Gal4}$ >$mCherry^{NLS}$ in $ap^{\Delta LSE}$ | *y w* ; $ap^{DG3}$ / $ap^{MS2}$<br>*y w* ; $ap^{DG3}$ / $ap^{MS2}$ ; $Dl^{Gal4}$ / *UAS*-$mCherry^{NLS}$ |
| **4d** | *ctrl*<br>$ap^{\Delta LSE}$ | *y w*<br>*y w* ; $ap^{DG3}$ / $ap^{MS2}$ |
| **5a-a´´´** | *ctrl*<br>$ci^{Rep}$<br>$byn^{Gal4}$<br>$byn^{Gal4}$ > $ci^{Rep}$ | *y w*<br>*y w* ; + ; *UAS*-$ci^{Rep}$ / +<br>*y w* ; + ; $byn^{Gal4}$ / +<br>*y w* ; + ; $byn^{Gal4}$ / *UAS*-$ci^{Rep}$ |
| **5b-g** | *ctrl*<br>$byn^{Gal4}$<br>$byn^{Gal4}$ in $ap^{\Delta LSE}$<br>$byn^{Gal4}$ > $ci^{Rep}$<br>$byn^{Gal4}$ > $ci^{Rep}$ in $ap^{\Delta LSE}$ | *y w*<br>*y w* ; + ; $byn^{Gal4}$ / +<br>*y w* ; $ap^{DG3}$ / $ap^{MS2}$ ; $byn^{Gal4}$ / +<br>*y w* ; + ; $byn^{Gal4}$ / *UAS*-$ci^{Rep}$<br>*y w* ; $ap^{DG3}$ / $ap^{MS2}$ ; $byn^{Gal4}$ / *UAS*-$ci^{Rep}$ |
| **6a-f** | *ctrl* | *y w* |
| **Sup 2a-d** | *ctrl*<br>$ap^{\Delta LSE}$ | *y w*<br>*y w* ; $ap^{DG3}$ / $ap^{MS2}$ |
| **Sup 3a-b** | *ctrl*<br>$ap^{\Delta LSE}$ | *y w*<br>*y w* ; $ap^{DG3}$ / $ap^{MS2}$ |
| **Sup 4** | *ctrl*<br>$ap^{\Delta LSE}$ | *y w*<br>*y w* ; $ap^{DG3}$ / $ap^{MS2}$ |
| **Sup 5a-b** | *ctrl* | *y w* |
| **Sup 6** | *ctrl*<br>$ap^{\Delta LSE}$ | *y w*<br>*y w* ; $ap^{DG3}$ / $ap^{MS2}$ |
| **Sup 7a** | *ctrl*<br>$ap^{\Delta LSE}$<br>$ap^{minLSE}$ | *y w*<br>*y w* ; $ap^{DG3}$ / $ap^{MS2}$<br>*y w* ; $ap^{MS2+minLSE}$ / $ap^{MS2}$ |
| **Sup 7b** | $byn^{Gal4}$ in $ap^{\Delta LSE}$<br>$byn^{Gal4}$ > $ci^{Rep}$<br>$byn^{Gal4}$ > $ci^{Rep}$ *in* $ap^{\Delta LSE}$ | *y w* ; $ap^{DG3}$ / $ap^{MS2}$ ; $byn^{Gal4}$ / +<br>*y w* ; + ; $byn^{Gal4}$ / *UAS*-$ci^{Rep}$<br>*y w* ; $ap^{DG3}$ / $ap^{MS2}$ ; $byn^{Gal4}$ / *UAS*-$ci^{Rep}$ |
| **Sub 7c-d** | *ctrl*<br>$ap^{\Delta LSE}$ | *y w*<br>*y w* ; $ap^{DG3}$ / $ap^{MS2}$ |
| **Sup 8 a-b** | *ctrl*<br>$byn^{Gal4}$ > $ci^{Rep}$ | *y w*<br>*y w* ; + ; $byn^{Gal4}$ / *UAS*-$ci^{Rep}$ |
| **Sup 9a-b** | *ctrl*<br>$byn^{Gal4}$ > $ci^{Rep}$<br>$byn^{Gal4}$ > $ci^{Rep}$ *in* $ap^{\Delta LSE}$ | *y w*<br>*y w* ; + ; $byn^{Gal4}$ / *UAS*-$ci^{Rep}$<br>*y w* ; $ap^{DG3}$ / $ap^{MS2}$ ; $byn^{Gal4}$ / *UAS*-$ci^{Rep}$ |
| **Sup 10a-b** | *ctrl* | *y w* |
