## Supplementary Table 3 for "Intestinal control of feeding initiation in *Drosophila melanogaster*"

### Supplementary table 3: Temperature shift experiment with *$ap^{md544}$ P{tub-Gal80$^{ts}$} / $ap^{DG3}$ P{UAS-ap)*

| temperature regime | @18°C* const. | day 5 shift | day 6 shift | day 7 shift | day 8 shift | day 9 shift | day 10 shift | day 11 shift | day 12 shift | day 13 shift | day 14 shift | day 15 shift | @29°C* const. |
|---|---|---|---|---|---|---|---|---|---|---|---|---|---|
| age[1] | | 96-120 | 120-144 | 144-168 | 168-192 | 192-216 | 216-240 | 240-264 | 264-288 | 288-312 | 312-336 | 336-360 | |
| survival[2] | 8/265 | 10/10 | 12/12 | 12/12 | 20/20 | 13/14 | 17/18 | 16/16 | 14/20 | 1/19 | 1/18 | 0/14 | 409/414 |
| fertility[3] | sterile | fertile | fertile | fertile | fertile | fertile | fertile | fertile | fertile | sterile | sterile | sterile | fertile |
| wings[4] | strong | normal | normal | strong | strong | strong | strong | strong | strong | strong | strong | strong | normal |

[*] observations made for cultures grown constantly at 18 or 29°C show that genotype *$ap^{md544}$ P{tub-Gal80$^{ts}$} / $ap^{DG3}$ P{UAS-ap)* is appropriate for our experiment. At 18°C, Gal80$^{ts}$ is functional and inhibits activation of *UAS-ap* by Gal4. Hence, no ap protein is produced and the typical ap phenotypes observed for genotype *$ap^{md544}$/$ap^{DG3}$* flies are fully penetrant. In contrast, at 29°C, Gal4 produced by *$ap^{md544}$* can activate *UAS-ap* and all ap phenotypes are rescued.

[1] indicates the age of the animals at the time they were shifted from 18 to 29°C. For example, “day 5 shift” means that embryos were collected at 18°C for 24 hrs. Then, “day 5 shift” animals were aged for 96 hrs at 18 °C before they were shifted to 29°C. Thus, at the time of the temperature shift, “day 5 shift” animals were 96 to 120 hrs old.

[2] indicates the number of adult flies that survived >3 days relative to the number of flies present on day 0.

[3] adult flies were also checked for their fertility. “sterile” means that no eggs were laid. “fertile” indicates that larval progeny were abundant. Note that precocious adult death and sterility phenotypes correlate (see also Wilson 1981a).

[4] *$ap^{md544}$/$ap^{DG3}$* flies display a wing phenotype close to that of *$ap^{null}$* flies. “strong” indicates that flies developed such wings. “normal” indicates that flies had wings like wild-type flies. Note that “day 6 shift” flies still develop normal wings. When using temperature sensitive allele *$ap^{ts78j}$*, Wilson reported that the temperature sensitive period for wing development extended “from late second through early third instar” (Wilson 1981a). It thus appears as if our assay system depending on Gal4/Gal80ts/UAS-ap is slightly delayed.
