## Supplementary Table 4 for "Intestinal control of feeding initiation in *Drosophila melanogaster*"

### Supplementary table 4: Temperature shift experiment with *byn$^{Gal4}$ P{tub-Gal80$^{ts}$} / P{UAS-ci$^{Rep}$)*

| temperature regime | @18°C* const. | day 3 shift | day 5 shift | day 7 shift | day 9 shift | day 11 shift | day 12 shift | day 13 shift | @29°C* const. |
|---|---|---|---|---|---|---|---|---|---|
| age[1] | | 48-72 | 96-120 | 144-168 | 192-216 | 240-264 | 264-288 | 288-312 | |
| survival[2] | >95% | >95% | >95% | >95% | >95% | >95% | >95% | >95% | >95% |
| papillae / ampulla[3] | | | | | | | | | |
| no papillae | - | 2/5 | 2/4 | 2/4 | 3/4 | 1/4 | - | - | 4/4 |
| 1 tiny papilla | - | 3/5 | 2/4 | 2/4 | 1/4 | 3/4 | - | - | - |
| 3 papillae | - | - | - | - | - | - | 1/5 | 1/3 | - |
| 4 papillae | 3/3 | - | - | - | - | - | 4/5 | 2/3 | - |
| fertility[4] | fertile | fertile | fertile | fertile | fertile | fertile | fertile | fertile | fertile |

* observations made for cultures grown constantly at 18 or 29°C show that genotype *by$^{Gal4}$ P{tub-Gal80$^{ts}$} / P{UAS-ci$^{Rep}$)* is appropriate for our experiment. At 18°C, Gal80$^{ts}$ is functional and inhibits activation of *UAS-ci$^{Rep}$* by Gal4. Hence, no ci$^{Rep}$ protein is produced. Ampullae develop normally and contain 4 papillae. In contrast, at 29°C, Gal4 produced by *byn$^{Gal4}$* can activate *UAS-ci$^{Rep}$* and papilla-less ampullae are formed.

[1] indicates the age of the animals at the time they were shifted from 18 to 29°C. For example, “day 3 shift” means that embryos were collected at 18°C for 24 hrs. Then, “day 3 shift” animals were aged for 48 hrs at 18 °C before they were shifted to 29°C. Thus, at the time of the temperature shift, “day 3 shift” animals were 48 to 72 hrs old.

[2] since essentially all flies were surviving well, no quantitative analysis of survival was done. >95% indicates that survival is basically normal.

[3] hindguts of a small number of female flies per shift were dissected and immediately analyzed and documented by light-microscopy. The number of papillae per ampulla is indicated. An example of a “tiny” papilla is shown in 9B’’.

[4] adult flies were also checked for their fertility. “fertile” indicates that larval progeny were abundant.
